# Comparative and longitudinal analysis of axial and retinal biometry in prospective models of hyperopia

**DOI:** 10.1101/2022.05.07.491049

**Authors:** Jai Pinkney, Navdeep Gogna, Gayle B. Collin, Lisa Stone, Mark P. Krebs, Juergen K. Naggert, Patsy M. Nishina

## Abstract

**Purpose:** To quantify changes in axial and retinal biometry in aging hyperopic mouse models.

**Methods:** Fundus photographs and ocular biometric measurements from *Mfrp*^*rd6*^, *Prss56*^*glcr4*^, *Adipor1*^*tm1Dgen*^, *C1qtnf5*^*tm1.1(KOMP)Vlcg*^ and *Prss56*^*em2(IMPC)J*^ homozygotes and C57BL/6J control mice were ascertained longitudinally up to one year of age. Parameters including axial length (AL), central corneal thickness (CCT), anterior chamber depth (ACD), lens thickness (LT), outer nuclear layer thickness (ONLT), retinal thickness (RT), vitreous chamber depth (VCD) and posterior length (PL) were measured using Spectral Domain-Optical Coherence Tomography imaging. Mixed-model analysis of variance and factorial analysis of covariance, using body size as a covariate, followed by post-hoc analysis was performed to identify significant strain differences.

**Results:** Strain specific changes in axial and retinal biometry along with significant effects of age, sex and body size on AL were noted. *Mfrp*^*rd6*^, *Prss56*^*glcr4*^, *Adipor1*^*tm1Dgen*^ and *Prss56*^*em2(IMPC)J*^ homozygotes had significantly shorter AL than controls. While a comparable decrease in PL was observed in *Mfrp*^*rd6*^, *Prss56*^*glcr4*^, and *Adipor1*^*tm1Dgen*^ homozygotes, the decrease was attributed to changes in different posterior components from each mutant. *Mfrp*^*rd6*^ and *Adipor1*^*tm1Dgen*^ homozygotes developed regularly sized fundus spots across the ocular globe, which differed from the large bright spots seen in aged *Prss56*^*glcr4*^ and *Prss56*^*em2(IMPC)J*^ homozygotes. While ONLT of *C1qtnf5*^*tm1.1(KOMP)Vlcg*^ mice was less than controls, AL and fundus images appeared normal.

**Conclusions:** This study highlights differences in contributions of ocular components to AL among hyperopic mouse models with decreased AL. Understanding the mechanisms through which these proteins function, will help to elucidate their role in controlling ocular growth.

## Introduction

Uncorrected refractive error (RE) is one of the leading causes of vision impairment and blindness in the United States and throughout the world^[1,2]^. Hyperopia (farsightedness), a common form of RE, is a condition where light rays from a nearby object get focused by the lens and converge behind the retina. Patients with hyperopia are unable to focus on nearby objects while seeing distant objects clearly. Biometric studies have shown that axial length (AL) is a significant determinant of hyperopia^[3,4]^.

A number of factors including race^[5]^, ethnicity^[5-7]^, sex, age and height^[8-12]^, time spent outdoors^[9]^, education^[6]^, and maternal smoking during pregnancy^[5]^ are associated with the onset and severity of hyperopia and/or changes in AL. Twin studies suggest that genetic factors account for as much as 86% of the variability seen in hyperopic cases^[13]^. In the Beaver Dam Study, 78% of the variability in hyperopic cases was attributed to genetic factors^[14]^. Genes associated with RE or AL in hyperopic patients have been found in familial linkage, candidate gene association and genome-wide association (GWA) studies^[15,16]^. Thus far, in humans, six genes: *MFRP, PRSS56, MYRF, TMEM98, CRB1*, and *BEST1* have been implicated in high hyperopia^[17-19]^ (≥ + 5.50 D) and nanophthalmos^[16,17,20-26]^ (+8.00 to +25.00 D), a severe form of this condition^[16]^.

Considering the many confounding variables outlined above, genetic models that can be studied in controlled settings are essential to uncover mechanisms and pathways underlying RE. The identification of the molecular players involved in hyperopia may offer new therapeutic targets to delay or ameliorate disease phenotypes. The genetic similarity to humans, the ability to control environment and genetic background, and the fast reproduction and maturation rates, altogether make mice valuable research models to study disease processes in the eye and other organ systems^[27]^. Since retinal structure, physiology and biochemistry are closely related in the mouse and human, murine disruption of human ocular disease genes often recapitulates disease phenotypes observed in affected patients. Finally, sensitized phenotypic screens in mice provide a powerful approach to identify gene interactions that contribute to ocular disease^[28-33]^. Accordingly, mice are well-positioned for advancing our understanding of hyperopia, particularly regarding the genetic and molecular networks that contribute to this condition. A number of murine genetic models for human hyperopia have been described^[34-38]^.

In 2011, Nair et. al.^[39]^ described *Prss56*^*grm4*^ (also referred as *Prss56*^*glcr4*^), a chemically-induced strain with a splice donor mutation in *Prss56*. Homozygous *Prss56*^*glcr4*^ mice have an increased anterior chamber depth, shortened AL, decreased posterior segment length, and angle-closure glaucoma. Paylakhi *et. al*.^[40]^, have recently shown that in addition to decreased AL, these mice are hyperopic and have an increased anterior chamber depth, decreased vitreous chamber depth, and increased retinal thickness. Similar to *MFRP* mutations in humans, mouse models with a disruption in *Mfrp* present with a shorter AL compared to controls^[28,29,36]^. *Mfrp*^*rd6*^ mice also have white fundus spots, progressive photoreceptor degeneration, and abnormal electroretinogram (ERG) responses^[37,38]^. Interestingly, Soundararjan et. al.^[41]^ showed that *Prss56* mRNA levels are increased by 17-fold and 70-fold in *Mfrp*^*rd6*^ eyes at P14 and in adult mice, respectively, suggesting an interaction between these two genes. *Mfrp* occurs as a dicistronic message with *C1qtnf5*; in prokaryotes, dicistronic genes share function and regulation^[42,43]^. Although the late-onset retinal degeneration (L-ORD) phenotype in *C1qtnf5* Ser163Arg knock-in^[44,45]^ models has been studied extensively, the effect of disrupting *C1qtnf5* on AL has not previously been examined. Notably, yeast two hybrid, co-immunoprecipitation and antibody localization studies have shown that *Mfrp* and *C1qtn5* interact^[46,47]^, which raises the possibility that models with *C1qtnf5* disruption might result in similar phenotypes as those with *Mfrp* mutations. Rice and coworkers^[48]^ described *Adipor1*^*-/-*^ (also referred as *Adipor1*^*tm1Dgen*^) mice which exhibit fundus spotting, photoreceptor degeneration, and abnormal ERGs, retinal phenotypes similar to those observed in *Mfrp*^*rd6*^ mutants. Recent studies by Gogna *et al* have shown that *Adipor1*^*tm1Dgen*^ mice have a shortened AL similar to *Mfrp*^*rd6*^ mutants^[28]^.

Here, as a first step in establishing whether prospective hyperopic mouse models recapitulate human hyperopic features, we used optical coherence tomography to assess axial and retinal biometric parameters. In addition to established models of hyperopia, *Mfrp*^*rd6*^, *Prss56*^*glcr4*^ and *Adipor1*^*tm1Dgen*^, we extended our analysis to two novel models generated by the Knockout Mouse Phenotyping Program (KOMP^2^) at The Jackson Laboratory (JAX), *C1qtnf5*^*tm1.1(KOMP)Vlcg*^ and Prss56^*em2(IMPC)J*^. We also examined the effects of age, sex, body weight and body length on AL measurements and performed a comparative analysis of select axial and retinal components. Our study highlights the complexity of the contributions to components of AL and suggests that the models described here may be used to elucidate the mechanisms that govern them.

## Materials and methods

### Animals

Mice used in this study were B6.C3Ga-*Mfrp*^*rd6*^/J (*JAX*^*®*^ mice, stock 003684^[37]^), B6(Cg)-*Prss56*^*glcr4*^/SjPjn^[39]^, C57BL/6NJ-*Prss56*^*em2(IMPC)J*^*/*Mmjax (MMRRC, stock 043778), B6.129P2-*Adipor1*^*tm1Dgen*^/Mmnc (MMRRC, stock 011599-UNC^[49]^), C57BL/6NJ - *C1qtnf5*^*tm1.1(KOMP)Vlcg*^/MMucd (*JAX*^*®*^ KOMP mice, stock JR18562^[50]^)and C57BL/6J (*JAX*^*®*^ *mice*, stock 000664), and are herein referred to as *Mfrp*^*rd6*^, *Prss56*^*glcr4*^, *Prss56*^*em2(IMPC)J*^, *Adipor1*^*tm1Dgen*^, *C1qtnf5*^*tm1.1(KOMP)Vlcg*^ and B6J, respectively. Unless otherwise indicated, mice used for characterization were homozygous for mutant alleles. Mice were bred and maintained under standardized conditions of 12:12 light:dark in the Research Animal Facility at The Jackson Laboratory (JAX). Mice were provided with LabDiet® 5K52 and HCl-acidified water (pH 2.8–3.2) *ad libitum* and maintained in pressurized individual ventilation caging. Mice were regularly monitored to ensure a pathogen-free environment. All strains were either already on B6J genetic background or were backcrossed to B6J for at least five generations and did not carry the *rd8* mutation. Animal protocols were approved by the JAX Institutional Animal Care and Use Committee (IACUC, AUS99089) and complied with guidelines set forth by the ARVO Animal Statement for the Use of Animals in Ophthalmic and Vision Research.

### Genotyping

DNA was extracted from tail and ear notch tissue specimens using a standard NaOH extraction protocol^[28]^. *Mfrp*^*rd6*^, *Adipor1*^*tm1Dgen*^ and *C1qtnf5*^*tm1.1(KOMP)Vlcg*^ mice were genotyped by the JAX Transgenic Genotyping Service while *Prss56*^*glcr4*[39]^ and *Prss56*^*em2(IMPC)J*^ mice were genotyped using an allele-specific PCR assay. PCR primers are listed in Supplemental Table ST1.

### RNA isolation and quantitative real-time (qRT) PCR

RNA isolation from posterior eye cups including retina was performed as described previously^[51]^ using the RNeasy Micro Kit (QIAGEN). cDNA was synthesized by reverse transcription of RNA using Superscript IV First Strand cDNA Synthesis kit (Thermofisher) according to manufacturer’s protocol. Subsequently, qRT-PCR was performed using a previously established protocol^[51]^, using iTaq Universal SYBR Green SuperMix (Bio-Rad), gene-specific primers (Supplementary Table ST1) and CFX96 Real-Time PCR Detection system (Bio-Rad). The comparative CT method (ΔΔCT) was applied to calculate a relative fold change in gene transcripts and was quantified using 2−ΔΔCT with *Actb* as an internal calibrator.

### Spectral Domain-Optical Coherence Tomography (SD-OCT) imaging of ocular parameters

Ocular biometric measurements were performed using a Bioptigen UltraHigh-resolution (UHR) Envisu R2210 SD-OCT imaging system (Leica microsystems). The principle of OCT^[27,52]^ and the method used to obtain images have been previously described^[53,54]^. Briefly, animals were anesthetized using ketamine (80 mg/kg)/xylazine (16 mg/kg), eyes were dilated with 1% cyclopentolate (Akorn, Inc) and kept hydrated using Systane and GenTeal eye drops. Eyes were kept in focus using the reference arm and focus dial, and aligned by positioning the Purkinje image at the center of the pupil. Cross-sectional images were obtained using A-scans (single depth profile composed of time-gated reflections) and B-scans (frame composed of array of A-scans). According to the manufacturer, our SD-OCT system captures an image depth of 1.653 mm in an A-scan of 1024 pixels, providing an axial resolution of 1.614μm/pixel.

#### Ocular measurements

A number of ocular biometric parameters were measured including AL, central anterior chamber depth (ACD), central corneal thickness (CCT), lens thickness (LT), vitreous chamber depth (VCD), outer nuclear layer thickness (ONLT), and retinal thickness (RT) and posterior length (PL) as shown in Figure 1A-B. While most of the ocular parameters could be measured using a single measurement, for the measurement of AL, the size limitations of the instrument’s image window limited our ability to capture the entire AL in a single measurement. We used the previously published approach of image inversion such that both the anterior and posterior segments of eye can be overlapped and seen within the same image window^[52]^ as shown in Figure 1C. AL was measured by taking a sum of three measurements – from the cornea to the image fold (bottom end), from the RPE/BM to image fold (top end), and one full length of the image (calibrated at refractive index of 1.653). For the measurement of LT, the sum of ACD, CT and PL was subtracted from AL. Previous studies have alternately used the term VCD to define the region from the back of the lens to the vitreal surface of retina^[55]^ or to the RPE/BM^[56]^. Since differences in RT can affect overall thickness of the region from back of the lens to the RPE/BM, particularly in small eyes, we preferred to refer to VCD as the distance to the vitreal surface of retina. We also wanted to examine if and how individual components i.e., vitreous chamber depth, retinal thickness and ONL thickness, contributed to the overall change in the posterior region of the eye in each of the mutant strains. Since the term posterior chamber has been used to describe the region between iris and lens^[57]^, we used the term ‘posterior length’ to describe the region from the back of the lens to RPE/BM. OCT images were randomized with anonymity, and age and strain identifiers were removed before analysis. Subsequently, images were evaluated using the Fiji open-source platform^[58]^.

**Figure 1:**
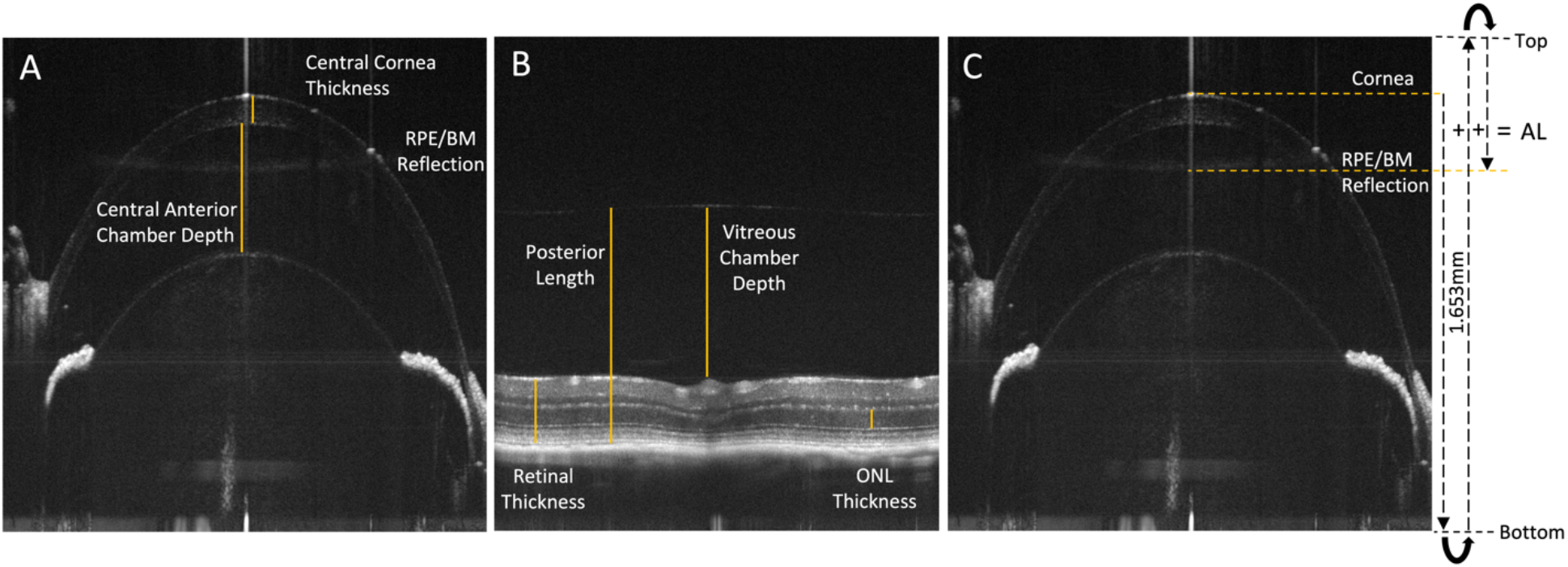
Representative *in vivo* OCT image of a C57BL/6J mouse. A. Ocular biometric parameters measured in the front of the eye including central anterior chamber depth (ACD) and central corneal thickness (CT) B. Ocular biometric parameters measured at the back of the eye including retinal thickness (RT), ONL thickness (ONLT), vitreous chamber depth (VCD), and posterior length (PL) C. Schematic diagram for the measurement of axial length, using the image inversion approach^[52]^, where the back of the eye can be seen as an inverted image in the same imaging window as the front of the eye. The axial length is measured as a sum of three measures – 1. measure from the front of the cornea to its inverting edge of the imaging window, 2. one full measure of the imaging window and 3. measure from the RPE/BM reflection to its inverting edge of the image window.

### Statistical Analysis

Data analysis was performed using JMP (SAS Institute, Cary, NC, USA) and GraphPad Prism (GraphPad Software, San Deigo, CA, USA) statistical analysis software. Mixed model analysis of variance (ANOVA), followed by Tukey’s multiple comparison post-hoc test was performed to identify significant differences in AL and other ocular parameters between different mouse strains at 1, 4, 8 and 12-months of age using GraphPad Prism. The results were validated by performing factorial ANOVA and three-way factorial analysis of covariance (ANCOVA) using the fit model function in JMP. AL was selected as a dependent variable, and main and interactive effects of three independent, categorical variables of mouse strains, age and gender were studied. Body size was used as a covariate. ANCOVA assumptions were tested before including body size as a covariate. A linear relationship between body size (covariate) and AL (dependent variable) as well as no interaction between the covariate and independent variables were confirmed. The data was also tested for normal distribution and equal variance. The F Ratio (ratio of the variation explained by the model and the unexplained variation; prob > F = < 0.05) and adjusted R^2^ (goodness-of-fit measure between the model and the dependent variable) and RMSE (Root Mean Square Error; measures the error of a model in predicting quantitative data and tells how accurately the model predicts the response; has the same unit as the quantity being measured) were obtained for the analysis and compared with and without the inclusion of covariate.

### Fundus imaging

Fundus photodocumentation was performed as previously described^[54]^, using a Micron IV fundus camera (Phoenix Research Laboratories, Pleasanton, CA, USA), with the exception that 1% cyclopentolate or 1% atropine was used as the dilating agent, and mice were anesthetized with isoflurane (isoflurane vaporizer from Kent Scientific, Torrington, CT, USA) during imaging.

## Results

### Strain selection for candidate models of hyperopia

To perform a detailed and comparative analysis of temporal changes in AL and retinal biometry, we selected aged *Mfrp*^*rd6*^, *Prss56*^*glcr4*^ and *Adipor1*^*tm1Dgen*^ murine models of genes which had previously been implicated in hyperopia and/or decreased AL^[28,36,40]^. In addition, we selected two uncharacterized JAX-KOMP^2^ strains; *Prss56*^*em2(IMPC)J*^, harboring a novel allele produced using Cas9-mediated genome editing^[59]^ and *C1qtnf5*^*tm1.1(KOMP)Vlcg*^, which harbors a reporter disruption in exons 2 and 3 of *C1qtnf5*. Given that *C1qtnf5* and *Mfrp* are expressed as dicistronic genes and are known to interact, we questioned whether this new knockout model would have phenotypic similarities with *Mfrp*^*rd6*^.

To assess the mutation effects on transcription levels in the two new KOMP-derived alleles, we performed quantitative RT-PCR on ocular cDNA specimens. *C1qtnf5* mRNA expression was significantly reduced in *C1qtnf5*^*tm1.1(KOMP)Vlcg*^ mutants compared to B6J controls (Supplementary Figure S1). The loss of mRNA may be attributed to nonsense mediated decay. Interestingly, *Prss56* mRNA expression was upregulated in *Prss56*^*em2(IMPC)J*^ homozygotes, as previously observed in *Prss56*^*glcr4*[39]^ mice (Supplementary Figure S2). The alleles and mutational effects on transcription and subsequent translated protein products are shown in Table 1.

**Table 1.** Candidate hyperopia models studied in this paper. *This paper; nd, not determined.

| Gene | Allele | Mutation | RNA | Protein |
| --- | --- | --- | --- | --- |
| *Adipor1* | *tm1Dgen* | Targeted deletion of c.107–198 in exon 2<sup>[49]</sup> | Reduced expression<sup>[60]</sup> | Loss of protein<sup>[49,60]</sup> |
| *C1qtnf5* | *tm1.1(KOMP)Vlcg* | Reporter-tagged deletion allele (1090 bp, part of exons 2 and 3) (MGI)<sup>[61,62]</sup> | Reduced expression* | nd |
| *Mfrp* | *rd6* | 4-bp deletion in intron 4 splice donor sequence resulting in a cryptic splice site causing in-frame skipping of exon 4<sup>[42]</sup> | Increased expression at P28<sup>[63]</sup> | Loss of protein<sup>[63]</sup> |
| *Prss56* | *glcr4* | T to A transversion in exon 11 splice donor sequence resulting in transcription of an additional 54 bp in intron 11 leading to a premature stop codon<sup>[39]</sup> | Increased expression<sup>[39]</sup> | nd |
| *Prss56* | *em2(IMPC)J* | Cas9-guided excision of 845 bp sequence removing exons 3 and 4 and 601 bp of flanking intron predicted to result in early termination due to loss of splice acceptor/donor(MGI)<sup>[61,62]</sup> | Increased expression* | nd |

### Comparative longitudinal analysis of AL

To determine the nature of axial changes in candidate mouse models of hyperopia (*Mfrp*^*rd6*^, *Prss56*^*glcr4*^ and *Adipor1*^*tm1Dgen*^, *Prss56*^*em2(IMPC)J*^ and *C1qtnf5*^*tm1.1(KOMP)Vlcg*^), AL measurements were ascertained and compared to B6J controls. Since AL changes with age^[64]^, comparative analysis was performed longitudinally over a period of one year to determine the onset and progression of axial changes. Initially, mixed-model ANOVA, followed by Tukey’s multiple comparison post-hoc analysis was used to compare AL values from different mouse strains at four timepoints (Figure 2). Significant main effects of mouse strain (F(DFn, DFd) = F(4, 100 = 70.03); *p*-value < 0.0001) and age (F(DFn, DFd) = F(2.769, 200.3 = 2560); *p*-value < 0.0001) were observed, confirming that AL differed significantly among mouse strains as well as across their range of ages. Significant interaction effects between strain and age (F(DFn, DFd) = F(12, 217 = 9.633); *p*-value < 0.0001) were also observed, suggesting that the axial differences between mouse strains were influenced by the age and vice versa. Adjusted *p* values obtained from Tukey’s multiple comparisons test for comparisons between multiple strains, at all ages, are listed in Supplementary Table ST2.

**Figure 2:**
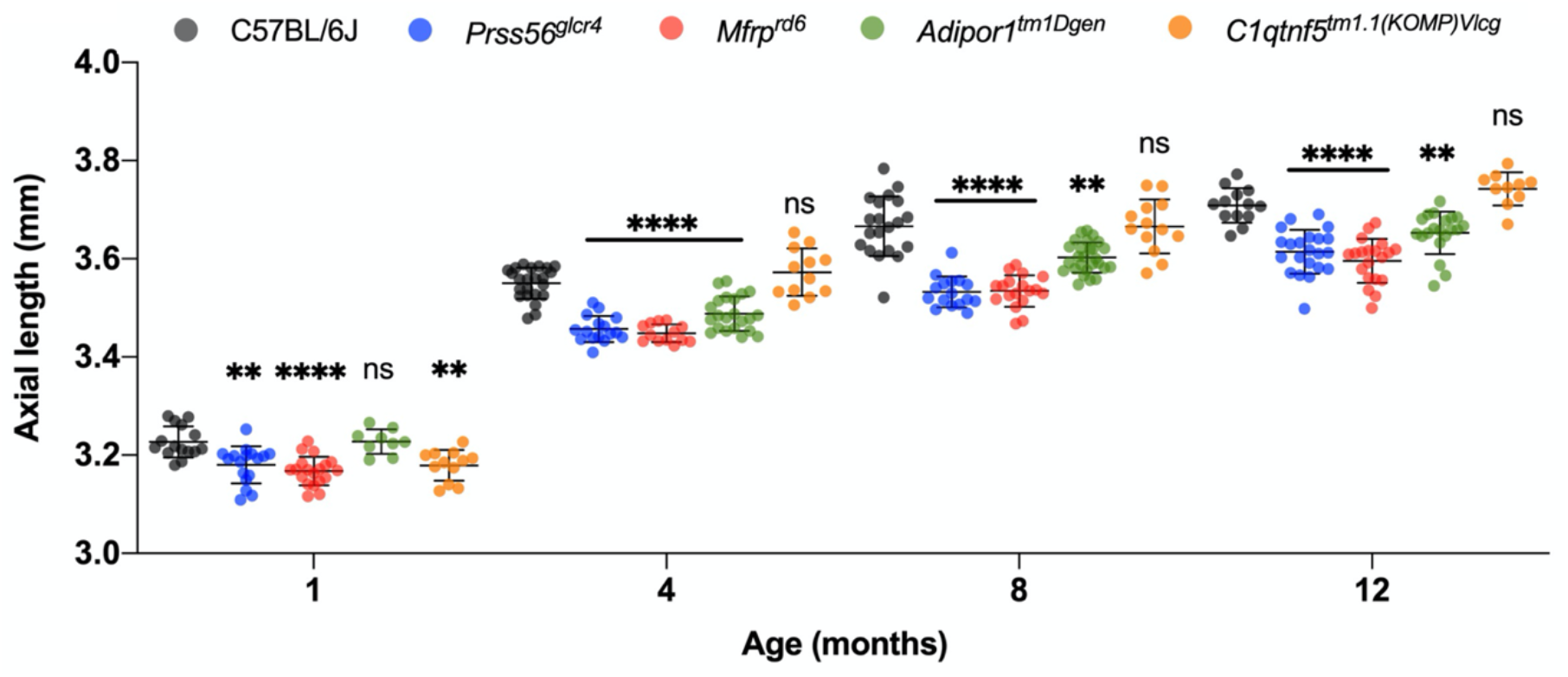
Comparative longitudinal analysis of AL in B6J control and prospective hyperopic mutant strains at 1, 4, 8 and 12 months of age. A progressive and significant decrease in AL was observed in *Prss56*^*glcr4*^, *Mfrp*^*rd6*^ and *Adipor1*^*tm1Dgen*^ homozygotes, whereas no change was detected in *C1qtnf5*^*tm1.1 (KOMP)Vlcg*^ homozygotes when compared to B6J controls, using mixed-model ANOVA, followed by Tukey’s multiple comparison test. B6J, *Prss56*^*glcr4*^, *Mfrp*^*rd6*^, *Adipor1*^*tm1Dgen*^ and *C1qtnf5*^*tm1.1(KOMP)Vlcg*^ mice are represented with gray, blue, red, green, and orange circles, respectively. Significant differences are indicated with asterisk (*) above each mutant with respect to B6J control animals. ** *p* < 0.01; **** *p* < 0.0001 and ns = not significant. n = 10-27 per strain per age, both sexes included.

### Effect of sex and body size on AL

To determine sex-based differences on AL, data was analyzed in JMP, where main and interaction effects of strain, age and sex were studied. An initial factorial ANOVA confirmed significant main and interaction effects for all three categorical variables (Table 2), suggesting that AL differed significantly among strains, between males and females, and at different ages. There was a significant main effect of sex, where AL was longer in males than females. Because males generally have larger body sizes than females, we performed ANCOVA and included body size as a covariate (Table 2) to establish whether the differences in AL may be due to variations in body size. Given that both body weight and body length are indicators of body size and are positively correlated to each other, we performed the dimension reduction method of principal component analysis (PCA) to obtain a single component representing body size. The ANCOVA analysis had a slightly better fit than the ANOVA analysis (ANOVA: adj *R*^*2*^ = 0.95, RMSE = 0.034 and F Ratio = 241.83 (Prob>F = <0.0001 vs ANCOVA: adj *R*^*2*^ = 0.97, RMSE = 0.031 and F Ratio = 253.84 (Prob>F = <0.0001). Thus, when controlling for body size, a significant but decreased effect of sex was observed, suggesting that body size contributed to the sex-based differences. The interaction terms involving sex were no longer significant after adjusting for body size, indicating that sex-based AL variations do not influence the differences due to strain and/or age. The main and interactive effects of strain and age were still significant, confirming that both strain and age were important factors leading to significant AL changes. Least squared means (LSMeans) differences, based on Tukey’s honestly significant differences (HSD) post-hoc analysis lists the significant differences between both the sexes of different mouse models at all four ages (Supplementary Table ST3).

**Table 2:** Effect tests showing main and interaction effects of strain, age and sex, with and without controlling for body size covariate. Significant F Ratios (Prob>F = <0.0001) are marked in red. After adjusting for covariate, the F Ratio for the sex term decreases and interaction terms involving sex become insignificant. The covariate significantly adjusted the dependent variable AL as determined from significant F Ratio (Prob>F = <0.0001).

| Factorial ANOVA |  |  | Factorial ANCOVA |  |  |
| --- | --- | --- | --- | --- | --- |
| Source | F Ratio | Prob > F | Source | F Ratio | Prob > F |
| Strain | 123.9277 | <0.0001 | Strain | 127.0379 | <0.0001 |
| Age | 2673.467 | <0.0001 | Age | 368.2881 | <0.0001 |
| Strain*Age | 10.7070 | <0.0001 | Strain*Age | 10.7204 | <0.0001 |
| Sex | 55.4749 | <0.0001 | Sex | 10.4014 | 0.0014 |
| Strain*Sex | 2.8913 | 0.0226 | Strain*Sex | 2.1440 | 0.0755 |
| Age*Sex | 0.8803 | 0.4517 | Age*Sex | 1.2268 | 0.3002 |
| Strain*Age*Sex | 1.8044 | 0.0469 | Strain*Age*Sex | 1.1228 | 0.3409 |
|  |  |  | Covariate (body size) | 22.7274 | <0.0001 |

### Longitudinal analysis of anterior axial measures

AL comprises several axial parameters. To determine how individual axial parameters vary among different mutant strains, we performed a comparative analysis for all the axial and retinal parameters, as shown in Figures 3 and 4 using mixed-model ANOVA followed by Tukey’s multiple comparison post-hoc test (Supplementary tables ST4-ST10). As seen in Figure 3, central corneal thickness (CT) did not differ significantly between mutants and control. While lens thickness (LT) increased longitudinally in all the strains, it did not differ significantly between most mutants and control. The anterior chamber depth (ACD) increased longitudinally in all the strains but differed with age such that all mutant strains, except *C1qtnf5*^*tm1.1(KOMP)Vlcg*^, had longer ACD than B6J controls.

**Figure 3:**
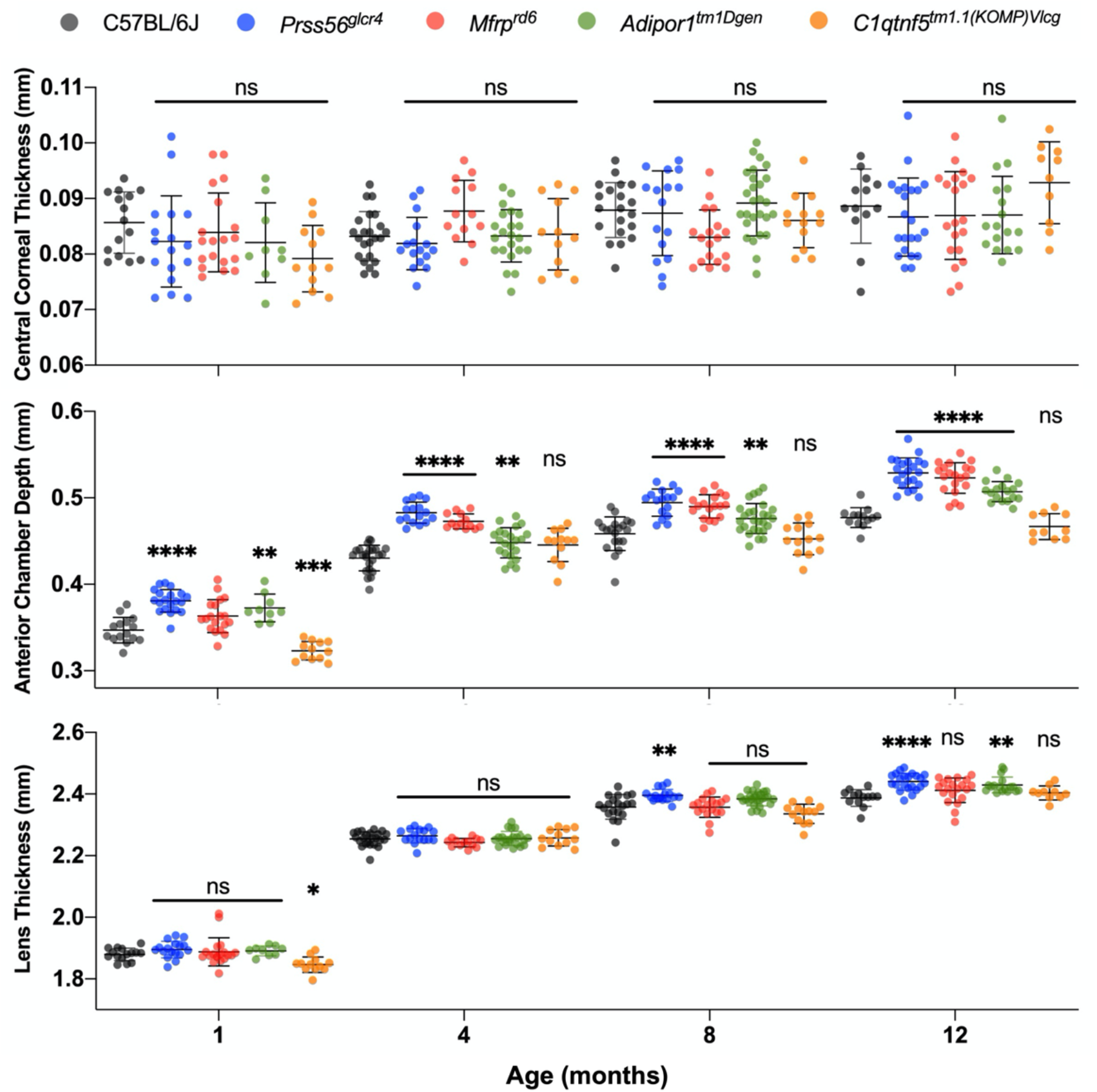
Comparative longitudinal analysis of ocular parameters in the anterior of the eye, for B6J control and mutant mice at 1, 4, 8 and 12 months of age. B6J, *Prss56*^*glcr4*^, *Mfrp*^*rd6*^, *Adipor1*^*tm1Dgen*^ and *C1qtnf5*^*tm1.1(KOMP)Vlcg*^ mice are represented in gray, blue, red, green and orange circles, respectively. Significant differences are indicated with asterisk (*) above each mutant with respect to B6J control animals. * *p* < 0.05; ** *p* < 0.01; *** *p* < 0.001; **** *p* < 0.0001 and ns = not significant. n = 10-27 per strain per age, both sexes included.

**Figure 4:**
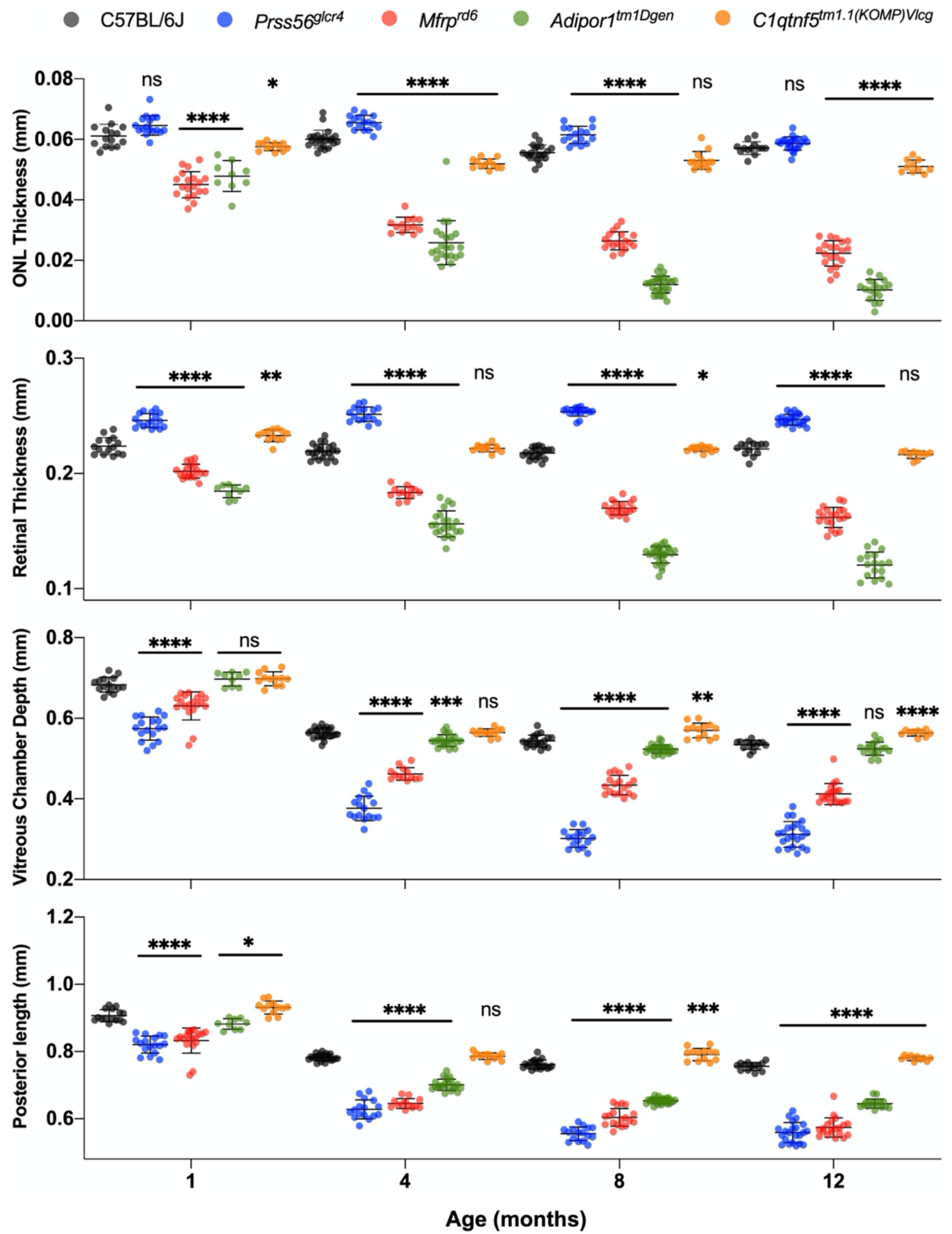
Comparative longitudinal analysis of ocular parameters in the posterior eye, for B6J control and mutant mice at 1, 4, 8 and 12 months of age. B6J, *Prss56*^*glcr4*^, *Mfrp*^*rd6*^, *Adipor1*^*tm1Dgen*^ and *C1qtnf5*^*tm1.1(KOMP)Vlcg*^ mice are represented with gray, blue, red, green, and orange circles, respectively. Significant differences are indicated with asterisk (*) above each mutant with respect to B6J control animals. * *p* < 0.05; ** *p* < 0.01; *** *p* < 0.001; **** *p* < 0.0001 and ns = not significant. n = 10-27 per strain per age, both sexes included.

### Longitudinal analysis of posterior axial measures

All posterior measurements including outer nuclear layer thickness (ONLT), retinal thickness (RT), vitreous chamber depth (VCD) and overall posterior length (PL) showed significant but different directions of change among the strains. The PL and VCD decreased both longitudinally and to a greater degree in the mutants as compared to control. The ONLT and RT decreased longitudinally and more significantly in *Mfrp*^*rd6*^ and *Adipor1*^*tm1Dgen*^ mice but not in *Prss56*^*glcr4*^ mice. Instead, there was an increase in ONLT and RT in *Prss56*^*glcr4*^ mice as the animals aged. Homozygous *C1qtnf5*^*tm1.1(KOMP)Vlcg*^ mice exhibited a slight increase in PL and VCD and a slight decrease in ONLT and RT compared to B6J controls.

### Time-course of fundus pathology by ophthalmoscopic imaging

*Mfrp*^*rd6*^ and *Adipor1*^*tm1Dgen*^ homozygotes share multiple disease phenotypes, including an early development of fundus spots^[28]^. To evaluate whether other strains present with abnormal fundus features, we compared strains longitudinally up to 12 months of age using ophthalmoscopic photodocumentation. (Figure 5). At one month, regularly sized, white fundus spots were detected in both *Mfrp*^*rd6*^ and *Adipor1*^*tm1Dgen*^ homozygotes and by four months, these spots became uniformly distributed across the ocular globe. In the fundus of *Prss56*^*glcr4*^ mice aged eight months and older, large, bright spots were found distributed along the central fundus and surrounded the optic nerve. Fundus phenotypes in *Prss56*^*em2(IMPC)J*^ homozygotes recapitulated those observed in *Prss56*^*glcr4*^ mice (Supplementary Figure S2). *C1qtnf5*^*tm1.1(KOMP)Vlcg*^ mutants had a normal fundus appearance throughout the 12-month time course.

**Figure 5:**
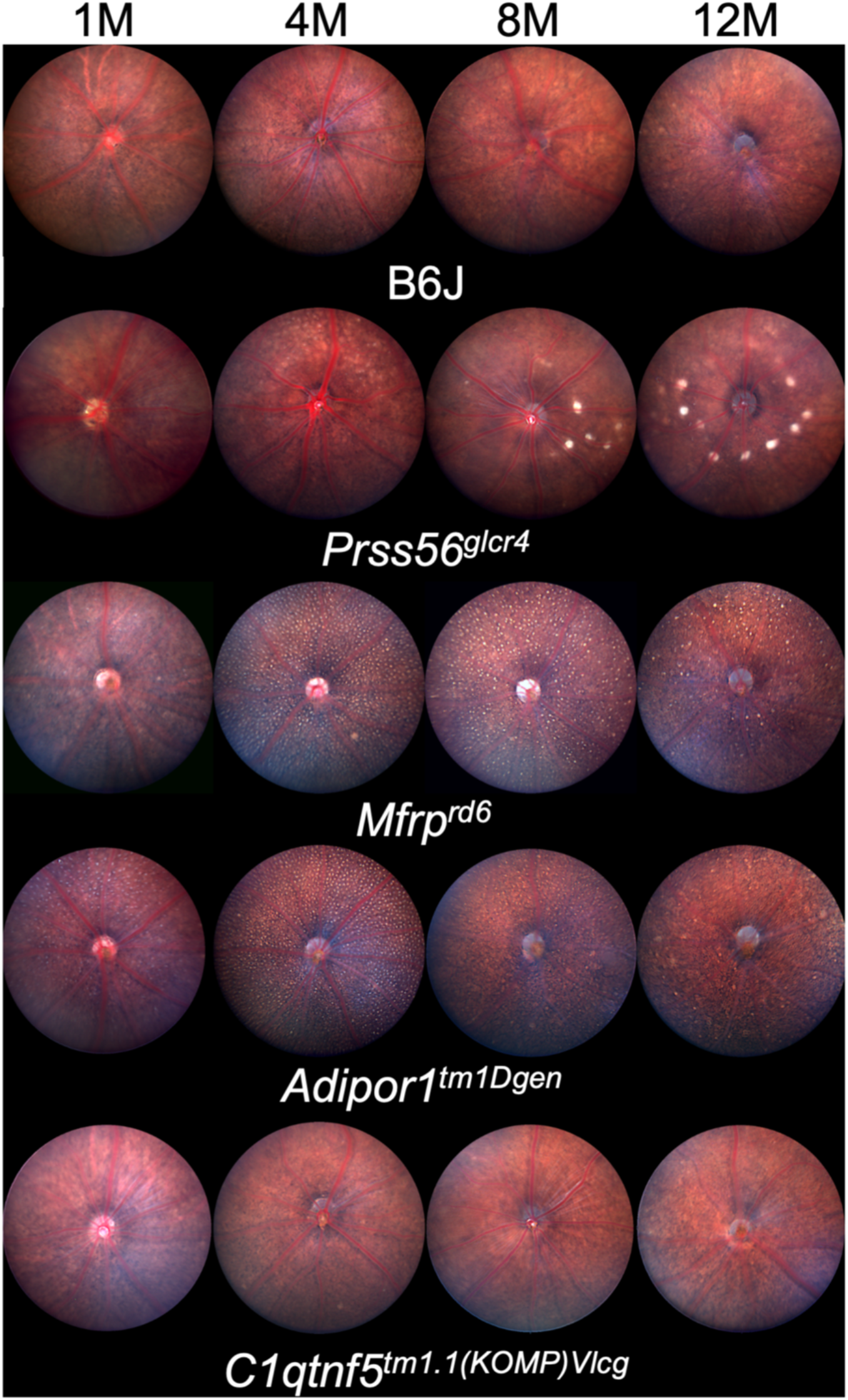
Fundus photographs of *Prss56*^*glcr4*^, *Mfrp*^*rd6*^, *Adipor1*^*tm1Dgen*^ and *C1qtnf5*^*tm1.1(KOMP)Vlcg*^ homozygotes and B6J controls, at 1, 4, 8 and 12 months of age, showing a progressive change in fundus appearance with age. While the uniformly sized spots in *Mfrp*^*rd6*^ and *Adipor1*^*tm1Dgen*^ mutants are similar in appearance and distribution, aged *Prss56*^*glcr4*^ mice develop large bright central fundus spots surrounding the optic nerve. *C1qtnf5*^*tm1.1(KOMP)Vlcg*^ mice do not develop spots and appear normal at all ages studied (n = 6-10, both sexes included/strain).

## Discussion

The World Health Organization estimates that greater than 2.2 billion people around the world are visually impaired, many of whom have uncorrected refractive errors such as hyperopia (WHO, accessed 10/14/2021). While AL contributes most significantly to hyperopia^[3]^, a few studies have implicated the contribution of other axial components in the development of hyperopia^[65]^. For example, a clinical study reported significantly lower ACD values in hyperopic patients^[66]^. In another correlation study, AL, ACD and VCD were strongly correlated with RE in hyperopic eyes^[67]^. A detailed understanding of how various compartments of the eye contribute to AL may help unravel the mechanisms that underlie the pathologic and molecular changes that result in hyperopia.

In order to understand how different hyperopic gene mutations affect the natural course of axial length changes, we examined the trajectory of AL and axial components longitudinally over 12 months. During this time course, we noted an early period of rapid AL growth, followed by a later period of slow growth (Figure 2). While some studies suggest that the mouse eye stops growing around postnatal day (P)40^[68]^, others suggest a two-phase growth pattern, with a period of rapid growth lasting until P40-P60, followed by a period of slow growth that continues up to P300^[64,69-71]^. The latter agrees with our findings; in all mutant strains studied, we observed an early period of rapid AL growth, followed by a slow growth stage. Moreover, our study confirmed mutant strain specific changes in AL and axial component’s growth trajectory in the mouse models *Mfrp*^*rd6*^, *Prss56*^*glcr4*^, *Adipor1*^*tm1Dgen*^ and *C1qtnf5*^*tm1.1(KOMP)Vlcg*^ (Figure 2-4). While an earlier and more significant AL change was observed in *Mfrp*^*rd6*^ and *Prss56*^*glcr4*^ homozygotes, the change progressed slower in *Adipor1*^*tm1Dgen*^ homozygotes. In our previous study, no change in AL was observed in heterozygous *Adipor1*^*tm1Dgen*^ mice at 4 months of age^[28]^. It is noteworthy that while heterozygous *Adipor1+/-* mice develop ocular phenotypes such as fundus spotting and PR degeneration, their progression is much slower than in homozygous *Adipor1-/-* mice. In our current study, we extended the analysis to 12 months of age to determine whether heterozygous animals develop a late AL phenotype. We did not observe any change in AL in heterozygous *Adipor1* mice at any of the timepoints examined (Supplementary Figure S3), indicating that decreased *Adipor1* levels affect disease phenotypes differently. Interestingly, while *C1qtnf5*^*tm1.1(KOMP)Vlcg*^ homozygotes had significantly shorter AL than controls at 1 month of age, the AL became comparable and insignificantly longer than the controls as the animals aged. Our results confirmed that although AL progressively increased with age in all the mice, the ALs of mutant strains were significantly shorter than the controls. Similar to previous reports^[11,72]^, we also observed significant differences between sexes for AL, which could be accounted for by including sex-based body size differences in the analysis.

In our comparative longitudinal analysis of axial components, we found that both PL and ACD contributed to overall changes in axial length. *Mfrp*^*rd6*^, *Prss56*^*glcr4*^, *Adipor1*^*tm1Dgen*^ mice showed a similar direction of change in AL and PL such that both decreased compared to control. On the other hand, we observed an opposite trend with ACD where mutants had increased size compared to controls. A shortening of the posterior segment is consistent with some human patients with disruptions in *MFRP* and *PRSS56*, which have been associated with posterior microphthalmia^[18,22,23,26,39,73-75]^. In mice, upon closer examination of the parameters that comprise PL, we noted that the reduction in PL results from decrease in different components of the eye in different mutants. For *Mfrp*^*rd6*^ mice, the decrease in PL could be attributed to a reduction in VCD, ONLT and RT. In the case of *Adipor1*^*tm1Dgen*^ mice, VCD did not show a dramatic change, and therefore, the smaller PL appeared to be due mainly to the decrease in RT and ONLT of the retina. For *Prss56*^*glcr4*^, the change in PL appeared to be mainly due to a smaller VCD. The RT and ONLT increased in *Prss56*^*glcr4*^. It is conceivable that the increase in RT and ONLT observed in *Prss56*^*glcr4*^ mice may be due to the fact that the amount of retina remains the same as in B6J but the area encompassing it decreases, leading to a compression of the retina in a smaller space. Interestingly, *Prss56*^*glcr4*^ homozygotes also present with retinal folds (Supplementary Figure 2), similar to the papillomacular folds observed in human *PRSS56* mutations that results from normal retinal development but arrested scleral growth^[18,23,39,73]^. The null *C1qtnf5*^*tm1.1(KOMP)Vlcg*^ mouse model showed decreased AL only at an early age of 1 month, and progressed towards increased AL thereafter. This increase in AL is accompanied by an increase in PL and VCD. However, it is important to note that ONLT was reduced in *C1qtnf5*^*tm1.1(KOMP)Vlcg*^ homozygotes, suggesting a slow degeneration phenotype in these mice.

Our comparative analysis of the fundus phenotype longitudinally at 1, 4, 8 and 12 months of age demonstrates key differences in ocular phenotypes between these disease models. While there was a progressive accumulation of fundus spots in *Mfrp*^*rd6*^, *Prss56*^*glcr4*^ and *Adipor1*^*tm1Dgen*^ mice with age, the spots differed in appearance and onset. *Mfrp*^*rd6*^ and *Adipor1*^*tm1Dgen*^ mutants developed uniformly sized and evenly distributed white spots across the fundus by 1-month of age, typical of the white flecks observed in human MFRP patients and in human flecked retinal disorder such as retinitis punctata albescens^[37,42]^. Our previous gene interaction study confirmed that *Mfrp* and *Adipor1* interact to affect the fundus spotting phenotype, suggesting a functional association between these genes^[28]^. The *Prss56*^*glcr4*^ mice on the other hand developed fundus spots that were bigger and brighter and appeared later (at 4 months) than the other two models. The retinal folds observed in *Prss56*^*glcr4*^ homozygotes appear to correspond to these fundus spots. AL measurements and fundus phenotypes of both the Prss56^*em2(IMPC)J*^ and *Prss56*^*glcr4*^ strains were comparable at the timepoints examined.

Throughout the twelve-month time course, *C1qtnf5*^*tm1.1(KOMP)Vlcg*^ mice did not develop a fundus phenotype and appeared normal. This contradicts previous reports where a *C1qtnf5* S163R knock-in mouse model (*C1qtnf5*^*tm1.1Itl*^) for late onset retinal degeneration (LORD), shows a fundus spotting phenotype^[45]^. However, the differences may be due to the fact that our model is a knock-out, confirmed by the lack of expression in qRT-PCR, whereas *C1qtnf5* knock-in mice expresses C1QTNF5 S163R protein. It is possible that the presence of an altered *C1qtnf5* results in disease whereas its complete absence does not. Additionally, the fundus phenotype reported for the S163R knock-in *C1qtnf5* heterozygotes was observed at 20-21 months of age^[45]^, a timepoint not assessed here. It remains possible that *C1qtnf5*^*tm1.1(KOMP)Vlcg*^ may have a late phenotype as well.

Our study highlights the underlying differences that contribute to decreased AL among different hyperopic models which share a smaller AL phenotype. Our study also prompts an interest to investigate the complex associations between *Mfrp* and related genes. It is conceivable for *Mfrp* to be associated with *Prss56* to affect the growth of vitreous chamber, and to be associated with *Adipor1* to control ONL and retinal growth, with both associations leading to alterations in posterior length. Unraveling the underlying mechanisms by which MFRP and other proteins act, will ultimately lead to a better understanding of how the proteins, when disrupted, results in RE and other ocular phenotypes.

## Acknowledgements

The authors thank Dr. Simon John for providing the *Prss56*^*glcr4*^ mice, Keith Sheppard for facilitating access to OCT data, Transgenic Genotyping Services for genotyping mice strains, and Melissa Berry for assistance with nomenclature. This research was funded by the National Eye Institute (NEI) grant, EY011996. The content is solely the responsibility of the authors and does not necessarily represent the official views of the National Institutes of Health. Core services at The Jackson Laboratory were supported by an Institutional grant, CA34196.

## References

1. Varma, R. et al. Visual Impairment and Blindness in Adults in the United States: Demographic and Geographic Variations From 2015 to 2050. JAMA Ophthalmol 134, 802–9 (2016).

2. Flaxman, S.R. et al. Global causes of blindness and distance vision impairment 1990-2020: a systematic review and meta-analysis. Lancet Glob Health 5, e1221–e1234 (2017).

3. Strang, N.C., Schmid, K.L. & Carney, L.G. Hyperopia is predominantly axial in nature. Curr Eye Res 17, 380–3 (1998).

4. Llorente, L., Barbero, S., Cano, D., Dorronsoro, C. & Marcos, S. Myopic versus hyperopic eyes: axial length, corneal shape and optical aberrations. J Vis 4, 288–98 (2004).

5. Borchert, M.S. et al. Risk factors for hyperopia and myopia in preschool children the multi-ethnic pediatric eye disease and Baltimore pediatric eye disease studies. Ophthalmology 118, 1966–73 (2011).

6. Wang, M. et al. Prevalence of and risk factors for refractive error: a cross-sectional study in Han and Mongolian adults aged 40-80 years in Inner Mongolia, China. Eye (Lond) 33, 1722–1732 (2019).

7. Ip, J.M. et al. Ethnic differences in refraction and ocular biometry in a population-based sample of 11-15-year-old Australian children. Eye (Lond) 22, 649–56 (2008).

8. Hashemi, H., Fotouhi, A. & Mohammad, K. The age- and gender-specific prevalences of refractive errors in Tehran: the Tehran Eye Study. Ophthalmic Epidemiol 11, 213–25 (2004).

9. Joseph, S. et al. Prevalence and risk factors for myopia and other refractive errors in an adult population in southern India. Ophthalmic Physiol Opt 38, 346–358 (2018).

10. Castagno, V.D., Fassa, A.G., Vilela, M.A., Meucci, R.D. & Resende, D.P. Moderate hyperopia prevalence and associated factors among elementary school students. Cien Saude Colet 20, 1449–58 (2015).

11. Kearney, S., Strang, N.C., Cagnolati, B. & Gray, L.S. Change in body height, axial length and refractive status over a four-year period in caucasian children and young adults. J Optom 13, 128–136 (2020).

12. Song, H.T., Kim, Y.J., Lee, S.J. & Moon, Y.S. Relations between age, weight, refractive error and eye shape by computerized tomography in children. Korean J Ophthalmol 21, 163–8 (2007).

13. Hammond, C.J., Snieder, H., Gilbert, C.E. & Spector, T.D. Genes and environment in refractive error: the twin eye study. Invest Ophthalmol Vis Sci 42, 1232–6 (2001).

14. Klein, A.P. et al. Heritability analysis of spherical equivalent, axial length, corneal curvature, and anterior chamber depth in the Beaver Dam Eye Study. Arch Ophthalmol 127, 649–55 (2009).

15. Young, T.L., Metlapally, R. & Shay, A.E. Complex trait genetics of refractive error. Arch Ophthalmol 125, 38–48 (2007).

16. Prasov, L. et al. Novel TMEM98, MFRP, PRSS56 variants in a large United States high hyperopia and nanophthalmos cohort. Sci Rep 10, 19986 (2020).

17. Orr, A. et al. Mutations in a novel serine protease PRSS56 in families with nanophthalmos. Mol Vis 17, 1850–61 (2011).

18. Gal, A. et al. Autosomal-recessive posterior microphthalmos is caused by mutations in PRSS56, a gene encoding a trypsin-like serine protease. Am J Hum Genet 88, 382–90 (2011).

19. Jiang, D. et al. Evaluation of PRSS56 in Chinese subjects with high hyperopia or primary angle-closure glaucoma. Mol Vis 19, 2217–26 (2013).

20. Lang, E. et al. Genotype-phenotype spectrum in isolated and syndromic nanophthalmos. Acta Ophthalmol 99, e594–e607 (2021).

21. Dudakova, L. et al. Pseudodominant Nanophthalmos in a Roma Family Caused by a Novel PRSS56 Variant. J Ophthalmol 2020, 6807809 (2020).

22. Siggs, O.M. et al. The genetic and clinical landscape of nanophthalmos and posterior microphthalmos in an Australian cohort. Clin Genet 97, 764–769 (2020).

23. Almoallem, B. et al. The majority of autosomal recessive nanophthalmos and posterior microphthalmia can be attributed to biallelic sequence and structural variants in MFRP and PRSS56. Sci Rep 10, 1289 (2020).

24. Guo, C. et al. Detection of Clinically Relevant Genetic Variants in Chinese Patients With Nanophthalmos by Trio-Based Whole-Genome Sequencing Study. Invest Ophthalmol Vis Sci 60, 2904–2913 (2019).

25. Carricondo, P.C., Andrade, T., Prasov, L., Ayres, B.M. & Moroi, S.E. Nanophthalmos: A Review of the Clinical Spectrum and Genetics. J Ophthalmol 2018, 2735465 (2018).

26. Said, M.B. et al. Posterior microphthalmia and nanophthalmia in Tunisia caused by a founder c.1059_1066insC mutation of the PRSS56 gene. Gene 528, 288–94 (2013).

27. Pardue, M.T., Stone, R.A. & Iuvone, P.M. Investigating mechanisms of myopia in mice. Exp Eye Res 114, 96–105 (2013).

28. Gogna, N. et al. Genetic Interaction between Mfrp and Adipor1 Mutations Affect Retinal Disease Phenotypes. Int J Mol Sci 23(2022).

29. Koli, S., Labelle-Dumais, C., Zhao, Y., Paylakhi, S. & Nair, K.S. Identification of MFRP and the secreted serine proteases PRSS56 and ADAMTS19 as part of a molecular network involved in ocular growth regulation. PLoS Genet 17, e1009458 (2021).

30. Maddox, D.M. et al. An allele of microtubule-associated protein 1A (Mtap1a) reduces photoreceptor degeneration in Tulp1 and Tub Mutant Mice. Invest Ophthalmol Vis Sci 53, 1663–9 (2012).

31. Li, S. et al. Nr2e3 is a genetic modifier that rescues retinal degeneration and promotes homeostasis in multiple models of retinitis pigmentosa. Gene Ther 28, 223–241 (2021).

32. Rao, K.N. et al. Ciliopathy-associated protein CEP290 modifies the severity of retinal degeneration due to loss of RPGR. Hum Mol Genet 25, 2005–2012 (2016).

33. Louie, C.M. et al. AHI1 is required for photoreceptor outer segment development and is a modifier for retinal degeneration in nephronophthisis. Nat Genet 42, 175–80 (2010).

34. Cross, S.H. et al. Missense Mutations in the Human Nanophthalmos Gene TMEM98 Cause Retinal Defects in the Mouse. Invest Ophthalmol Vis Sci 60, 2875–2887 (2019).

35. Garnai, S.J. et al. Variants in myelin regulatory factor (MYRF) cause autosomal dominant and syndromic nanophthalmos in humans and retinal degeneration in mice. PLoS Genet 15, e1008130 (2019).

36. Velez, G. et al. Gene Therapy Restores Mfrp and Corrects Axial Eye Length. Sci Rep 7, 16151 (2017).

37. Hawes, N.L. et al. Retinal degeneration 6 (rd6): a new mouse model for human retinitis punctata albescens. Invest Ophthalmol Vis Sci 41, 3149–57 (2000).

38. Fogerty, J. & Besharse, J.C. 174delG mutation in mouse MFRP causes photoreceptor degeneration and RPE atrophy. Invest Ophthalmol Vis Sci 52, 7256–66 (2011).

39. Nair, K.S. et al. Alteration of the serine protease PRSS56 causes angle-closure glaucoma in mice and posterior microphthalmia in humans and mice. Nat Genet 43, 579–84 (2011).

40. Paylakhi, S. et al. Muller glia-derived PRSS56 is required to sustain ocular axial growth and prevent refractive error. PLoS Genet 14, e1007244 (2018).

41. Soundararajan, R. et al. Gene profiling of postnatal Mfrprd6 mutant eyes reveals differential accumulation of Prss56, visual cycle and phototransduction mRNAs. PLoS One 9, e110299 (2014).

42. Kameya, S. et al. Mfrp, a gene encoding a frizzled related protein, is mutated in the mouse retinal degeneration 6. Hum Mol Genet 11, 1879–86 (2002).

43. Chavali, V.R., Sommer, J.R., Petters, R.M. & Ayyagari, R. Identification of a promoter for the human C1Q-tumor necrosis factor-related protein-5 gene associated with late-onset retinal degeneration. Invest Ophthalmol Vis Sci 51, 5499–507 (2010).

44. Shu, X. et al. Characterisation of a C1qtnf5 Ser163Arg knock-in mouse model of late-onset retinal macular degeneration. PLoS One 6, e27433 (2011).

45. Chavali, V.R. et al. A CTRP5 gene S163R mutation knock-in mouse model for late-onset retinal degeneration. Hum Mol Genet 20, 2000–14 (2011).

46. Shu, X. et al. Disease mechanisms in late-onset retinal macular degeneration associated with mutation in C1QTNF5. Hum Mol Genet 15, 1680–9 (2006).

47. Mandal, M.N. et al. Spatial and temporal expression of MFRP and its interaction with CTRP5. Invest Ophthalmol Vis Sci 47, 5514–21 (2006).

48. Rice, D.S. et al. Adiponectin receptor 1 conserves docosahexaenoic acid and promotes photoreceptor cell survival. Nat Commun 6, 6228 (2015).

49. Bjursell, M. et al. Opposing effects of adiponectin receptors 1 and 2 on energy metabolism. Diabetes 56, 583–93 (2007).

50. Sundberg, J.P. et al. Systematic screening for skin, hair, and nail abnormalities in a large-scale knockout mouse program. PLoS One 12, e0180682 (2017).

51. Collin, G.B. et al. Disruption of murine Adamtsl4 results in zonular fiber detachment from the lens and in retinal pigment epithelium dedifferentiation. Hum Mol Genet 24, 6958–74 (2015).

52. Park, H. et al. Assessment of axial length measurements in mouse eyes. Optom Vis Sci 89, 296–303 (2012).

53. Krebs, M.P. Using Vascular Landmarks to Orient 3D Optical Coherence Tomography Images of the Mouse Eye. Curr Protoc Mouse Biol 7, 176–190 (2017).

54. Krebs, M.P., Xiao, M., Sheppard, K., Hicks, W. & Nishina, P.M. Bright-Field Imaging and Optical Coherence Tomography of the Mouse Posterior Eye. Methods Mol Biol 1438, 395–415 (2016).

55. Chou, T.H. et al. Postnatal elongation of eye size in DBA/2J mice compared with C57BL/6J mice: in vivo analysis with whole-eye OCT. Invest Ophthalmol Vis Sci 52, 3604–12 (2011).

56. Takkar, B. et al. Evaluation of the vitreous chamber depth: An assessment of correlation with ocular biometrics. Indian J Ophthalmol 67, 1645–1649 (2019).

57. Choi, W.J., Pepple, K.L., Zhi, Z. & Wang, R.K. Optical coherence tomography based microangiography for quantitative monitoring of structural and vascular changes in a rat model of acute uveitis in vivo: a preliminary study. J Biomed Opt 20, 016015 (2015).

58. Schindelin, J. et al. Fiji: an open-source platform for biological-image analysis. Nat Methods 9, 676–82 (2012).

59. Elrick, H. et al. The production of 4,182 mouse lines identifies experimental and biological variables impacting Cas9-mediated mutant mouse line production.

60. Sluch, V.M. et al. ADIPOR1 is essential for vision and its RPE expression is lost in the Mfrp(rd6) mouse. Sci Rep 8, 14339 (2018).

61. Lloyd, K.C. A knockout mouse resource for the biomedical research community. Ann N Y Acad Sci 1245, 24–6 (2011).

62. Dickinson, M.E. et al. High-throughput discovery of novel developmental phenotypes. Nature 537, 508–514 (2016).

63. Won, J. et al. Membrane frizzled-related protein is necessary for the normal development and maintenance of photoreceptor outer segments. Vis Neurosci 25, 563–74 (2008).

64. Zhou, G. & Williams, R.W. Mouse models for the analysis of myopia: an analysis of variation in eye size of adult mice. Optom Vis Sci 76, 408–18 (1999).

65. Uretmen, O., Pamukcu, K., Kose, S. & Egrilmez, S. Oculometric features of hyperopia in children with accommodative refractive esotropia. Acta Ophthalmol Scand 81, 260–3 (2003).

66. Rabsilber, T.M., Becker, K.A., Frisch, I.B. & Auffarth, G.U. Anterior chamber depth in relation to refractive status measured with the Orbscan II Topography System. J Cataract Refract Surg 29, 2115–21 (2003).

67. Chang, C.K., Lin, J.T. & Zhang, Y. Correlation analysis and multiple regression formulas of refractive errors and ocular components. Int J Ophthalmol 12, 858–861 (2019).

68. Schmucker, C. & Schaeffel, F. In vivo biometry in the mouse eye with low coherence interferometry. Vision Res 44, 2445–56 (2004).

69. Zhou, X. et al. The development of the refractive status and ocular growth in C57BL/6 mice. Invest Ophthalmol Vis Sci 49, 5208–14 (2008).

70. Puk, O., Dalke, C., Favor, J., de Angelis, M.H. & Graw, J. Variations of eye size parameters among different strains of mice. Mamm Genome 17, 851–7 (2006).

71. Schmucker, C. & Schaeffel, F. A paraxial schematic eye model for the growing C57BL/6 mouse. Vision Res 44, 1857–67 (2004).

72. Roy, A., Kar, M., Mandal, D., Ray, R.S. & Kar, C. Variation of Axial Ocular Dimensions with Age, Sex, Height, BMI-and Their Relation to Refractive Status. J Clin Diagn Res 9, AC01–4 (2015).

73. Sen, P. et al. Long-term follow-up of a case of posterior microphthalmos (PRSS56) with hyperautofluorescent retinal pigment epithelial deposits. Eur J Ophthalmol 32, NP163–NP167 (2022).

74. Bacci, G.M. et al. Novel mutations in MFRP and PRSS56 are associated with posterior microphthalmos. Ophthalmic Genet 41, 49–56 (2020).

75. Nowilaty, S.R. et al. Biometric and molecular characterization of clinically diagnosed posterior microphthalmos. Am J Ophthalmol 155, 361–372 e7 (2013).

