## Supplementary for "Comparative and longitudinal analysis of axial and retinal biometry in prospective models of hyperopia"

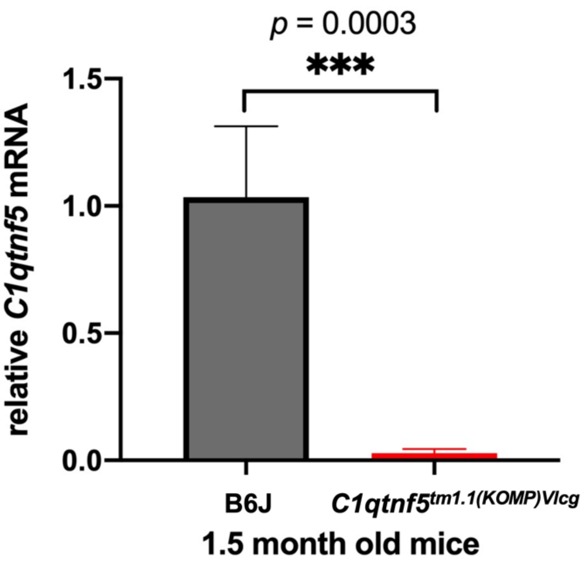


**Figure S1:** Expression analysis of *C1qtnf5*$^{tm1.1(KOMP)Vlcg}$ by qRT-PCR. A significant reduction of *C1qtnf5* expression was observed in posterior eyecups from 1.5-month-old mutant mice when compared to B6J controls, n = 6/strain, both sexes included.

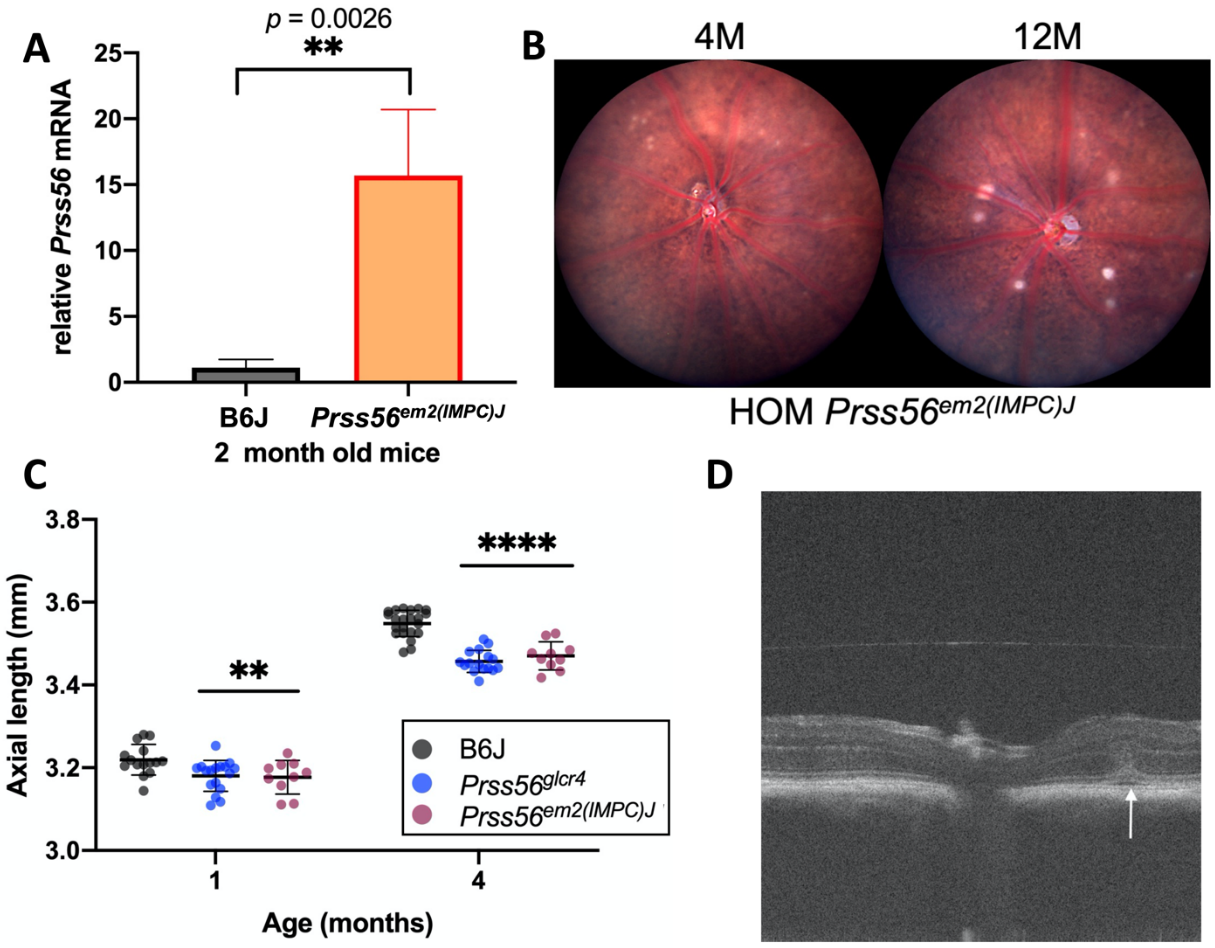


**Figure S2:** Molecular and phenotypic similiarites among *Prss56* allelic variants. **A.** qRT-PCR using two-month-old whole eye cDNA showed a significant upregulation of *Prss56* expression in $Prss56^{em2(IMPC)J}$ homozygotes compared to B6J controls, n = 4-5/strain **B.** Fundus photos showing bright spots in $Prss56^{em2(IMPC)J}$ homozygotes (HOM), similar to that observed in $Prss56^{glcr4}$ mice, at 4 and 12 months. n = 5/age **C.** Comparative AL analysis shows a similar decrease in AL in both $Prss56^{glcr4}$ and $Prss56^{em2(IMPC)J}$ mice at 1 and 4 months of age, respectively. n = 10-17 mice/strain, both sexes included. ** $p < 0.01$; **** $p < 0.0001$. **D**. SD-OCT image showing a retinal fold (arrow) in the posterior eye of a 12-month-old $Prss56^{glcr4}$ mouse, similar to the papillomacular folds seen in human *PRSS56* patients.

.

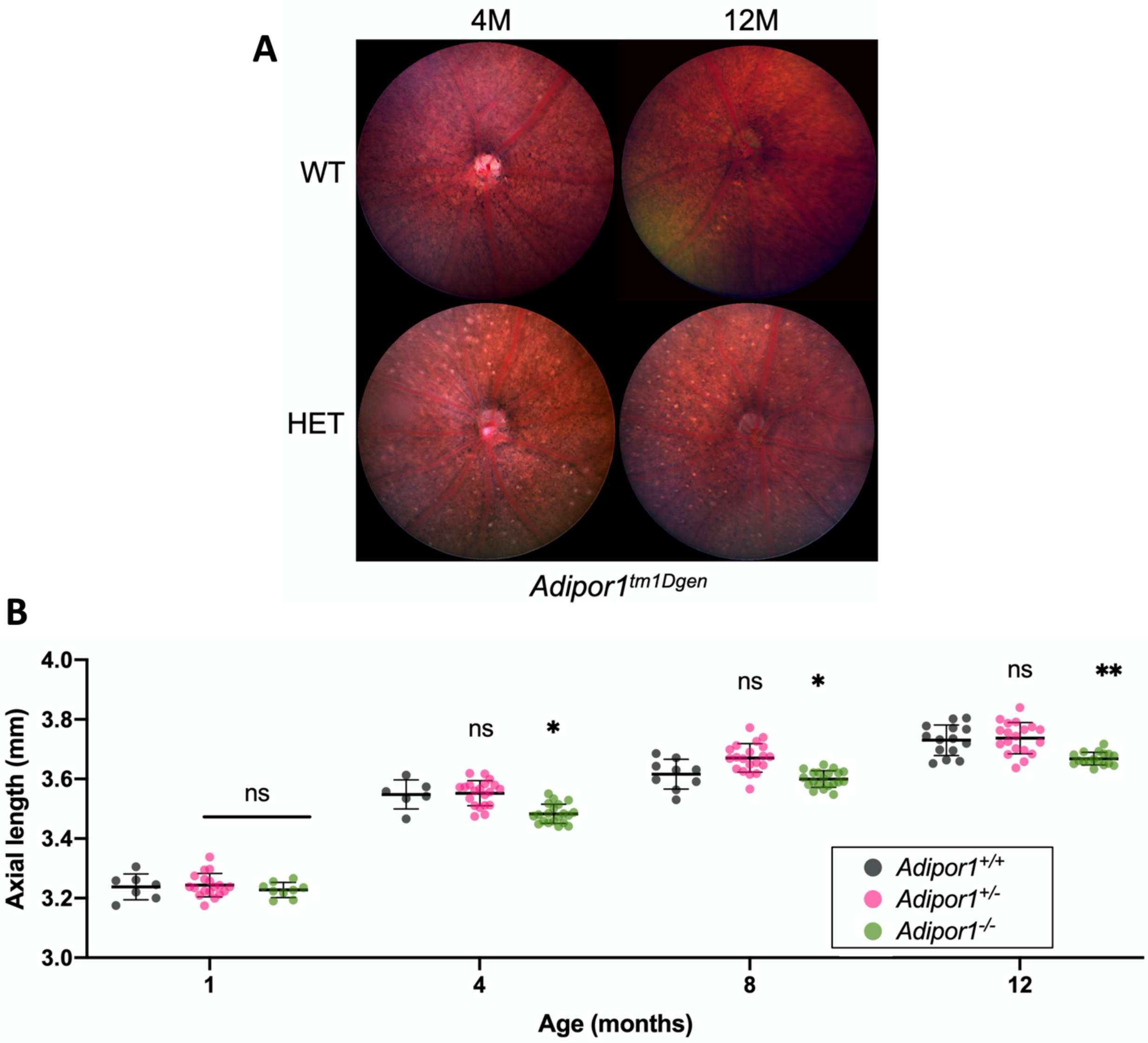


**Figure S3:** Effects of zygosity on ocular phenotypes in *Adipor1*$^{tm1Dgen}$ mice **A.** *top panel*, normal fundus of a wild-type (+/+) littermate control from an F2 cross segregating for *Adipor1* ko allele. *bottom panel,* fundus spotting in heterozygous (HET) *Adipor1*$^{tm1Dgen}$ mice, shown at 4 and 12 months of age n = 5/cohort **B.** Comparative longitudinal analysis shows that AL in heterozygous *Adipor1+/-* mice (pink circles) resembles wild-type controls (grey circles) whereas AL is reduced in homozygous *Adipor1 -/-* mice (green circles). * $p < 0.05$; ** $p < 0.01$, n = 10-20 mice/strain, both genders included.

| ***GENOTYPING*** | | | | | |
|---|---|---|---|---|---|
| **Strain** | **Allele** | **Size (bp)** | **Forward** | **Reverse** | **Assay** |
| *Adipor1*$^{tm1Dgen}$ | WT | 265 | TTA GAG GCA GGG TAA GCT GAT | AGC CAG CTC CAC TGT GTC AG | MCS |
| | MUT | ~450 | | TGG GAT TAG ATA AAT GCC TGC TCT | |
| *C1qtnf5*$^{tm1.1(KOMP)Vlcg}$ | WT | 382 | CTACCTTTCGACCGTGTGCT | CCGGGTTCACTGTGTTTTAAG | S |
| | MUT | 729 | CGG TCG CTA CCA TTA CCA GT | GGA GCA GCA GAG ATG GAG TC | |
| *Mfrp*$^{rd6}$ | WT | 116 | ACTACCACCCCAGCAAGGAC<br>Probe:5HEX-CAGACCAGTAAGTCCCAAGGG | CTTTCCTCCCCAACACCATC<br>Probe:6FAM-CAGACCAGTCCCAAGGGC | EP |
| | MUT | 112 | | | |
| *Prss56*$^{glcr4}$ | WT | 175 | TGGCTCCAGAAACCAAAGCCGGAAGAGCGCCCGGAAACAAAGAGT | TCCTGGAAGAGAGGGAGTGA (Common) | AS |
| | MUT | 152 | GCGGCGCCCGGAAACAAAAGGA | | |
| *Prss56*$^{em2(IMPC)J}$ | WT | 104 | CCCAGGGTAGGAGAACATCA | GCCTGTAAGTGTGGCTGTTG (Common) | S |
| | MUT | 120 | GGTTGTACCAGAAGTGCTACCC | | |

| ***qRT-PCR*** | | | |
|---|---|---|---|
| **Gene (primers)** | **Size (bp)** | **Forward** | **Reverse** |
| *C1qtnf5* (F1/R1) | 126 | GAGACCGGGACTACCTG | GATCGCTTGGCACTGAA |
| *C1qtnf5* (F2/R2) | 117 | GCACACCAGGTCACCAT | CAGGTAGTCCCGGTCTC |
| *Prss56* (F/1R) | 183 | CTGTGGACTGTGATGCTCGC | ACCCTGGGGAAGGCAAAT |
| *Actb* (F/R) | 207 | CCAGTTCGCCATGGATGACGATAT | GTCAGGATACCTCTCTTGCTCTG |

**Table ST1:** Primers used for genotyping and qRT-PCR. AS, Allele-specific,; EP, End-Point; MCS, Melting Curve; S, Standard PCR

| Tukey's multiple comparison | Adjusted p-value | Summary |
|---|---|---|
| **Age (1 month)** | | |
| C57BL/6J vs. $Mfrp^{rd6}$ | <0.0001 | **** |
| C57BL/6J vs. $Prss56^{glcr4}$ | 0.0056 | ** |
| C57BL/6J vs. $Adipor1^{tm1Dgen}$ | >0.9999 | ns |
| C57BL/6J vs. $C1qtnf5^{tm1.1(KOMP)Vlcg}$ | 0.0058 | ** |
| $Mfrp^{rd6}$ vs. $Prss56^{glcr4}$ | 0.7907 | ns |
| $Mfrp^{rd6}$ vs. $Adipor1^{tm1Dgen}$ | 0.0002 | *** |
| $Mfrp^{rd6}$ vs. $C1qtnf5^{tm1.1(KOMP)Vlcg}$ | 0.8281 | ns |
| $Prss56^{glcr4}$ vs. $Adipor1^{tm1Dgen}$ | 0.0074 | ** |
| $Prss56^{glcr4}$ vs. $C1qtnf5^{tm1.1(KOMP)Vlcg}$ | >0.9999 | ns |
| $Adipor1^{tm1Dgen}$ vs. $C1qtnf5^{tm1.1(KOMP)Vlcg}$ | 0.0075 | ** |
| **Age (4 month)** | | |
| C57BL/6J vs. $Mfrp^{rd6}$ | <0.0001 | **** |
| C57BL/6J vs. $Prss56^{glcr4}$ | <0.0001 | **** |
| C57BL/6J vs. $Adipor1^{tm1Dgen}$ | <0.0001 | **** |
| C57BL/6J vs. $C1qtnf5^{tm1.1(KOMP)Vlcg}$ | 0.6058 | ns |
| $Mfrp^{rd6}$ vs. $Prss56^{glcr4}$ | 0.8262 | ns |
| $Mfrp^{rd6}$ vs. $Adipor1^{tm1Dgen}$ | 0.0008 | *** |
| $Mfrp^{rd6}$ vs. $C1qtnf5^{tm1.1(KOMP)Vlcg}$ | <0.0001 | **** |
| $Prss56^{glcr4}$ vs. $Adipor1^{tm1Dgen}$ | 0.0274 | * |
| $Prss56^{glcr4}$ vs. $C1qtnf5^{tm1.1(KOMP)Vlcg}$ | <0.0001 | **** |
| $Adipor1^{tm1Dgen}$ vs. $C1qtnf5^{tm1.1(KOMP)Vlcg}$ | 0.0004 | *** |
| **Age (8 month)** | | |
| C57BL/6J vs. $Mfrp^{rd6}$ | <0.0001 | **** |
| C57BL/6J vs. $Prss56^{glcr4}$ | <0.0001 | **** |
| C57BL/6J vs. $Adipor1^{tm1Dgen}$ | 0.0019 | ** |
| C57BL/6J vs. $C1qtnf5^{tm1.1(KOMP)Vlcg}$ | >0.9999 | ns |
| $Mfrp^{rd6}$ vs. $Prss56^{glcr4}$ | 0.9998 | ns |
| $Mfrp^{rd6}$ vs. $Adipor1^{tm1Dgen}$ | <0.0001 | **** |
| $Mfrp^{rd6}$ vs. $C1qtnf5^{tm1.1(KOMP)Vlcg}$ | <0.0001 | **** |
| $Prss56^{glcr4}$ vs. $Adipor1^{tm1Dgen}$ | <0.0001 | **** |
| $Prss56^{glcr4}$ vs. $C1qtnf5^{tm1.1(KOMP)Vlcg}$ | <0.0001 | **** |
| $Adipor1^{tm1Dgen}$ vs. $C1qtnf5^{tm1.1(KOMP)Vlcg}$ | 0.0107 | * |
| **Age (12 month)** | | |
| C57BL/6J vs. $Mfrp^{rd6}$ | <0.0001 | **** |
| C57BL/6J vs. $Prss56^{glcr4}$ | <0.0001 | **** |
| C57BL/6J vs. $Adipor1^{tm1Dgen}$ | 0.0028 | ** |
| C57BL/6J vs. $C1qtnf5^{tm1.1(KOMP)Vlcg}$ | 0.1710 | ns |
| $Mfrp^{rd6}$ vs. $Prss56^{glcr4}$ | 0.6575 | ns |
| $Mfrp^{rd6}$ vs. $Adipor1^{tm1Dgen}$ | 0.0018 | ** |
| $Mfrp^{rd6}$ vs. $C1qtnf5^{tm1.1(KOMP)Vlcg}$ | <0.0001 | **** |
| $Prss56^{glcr4}$ vs. $Adipor1^{tm1Dgen}$ | 0.0526 | ns |
| $Prss56^{glcr4}$ vs. $C1qtnf5^{tm1.1(KOMP)Vlcg}$ | <0.0001 | **** |
| $Adipor1^{tm1Dgen}$ vs. $C1qtnf5^{tm1.1(KOMP)Vlcg}$ | <0.0001 | **** |

**Table ST2:** Adjusted *p*-values obtained from Tukey's multiple comparison test for AL comparison between different mouse strains, at four different ages.

| Strain | Age | Sex | Level | Least Sq Mean |
|---|---|---|---|---|
| $C1qtnf5^{tm1.1(KOMP)Vlcg}$ | 12 | M | A | 3.7297960 |
| $C1qtnf5^{tm1.1(KOMP)Vlcg}$ | 12 | F | A,B | 3.7091270 |
| B6J | 12 | M | A,B,C | 3.6913368 |
| B6J | 8 | M | A,B,C | 3.6787285 |
| $C1qtnf5^{tm1.1(KOMP)Vlcg}$ | 8 | F | A,B,C,D | 3.6662968 |
| B6J | 12 | F | A,B,C,D | 3.6606941 |
| $C1qtnf5^{tm1.1(KOMP)Vlcg}$ | 8 | M | A,B,C,D,E | 3.6523358 |
| *Adipor1* | 12 | M | B,C,D,E,F | 3.6381963 |
| *B6J* | 8 | F | C,D,E,F,G | 3.6343285 |
| *Adipor1* | 12 | F | C,D,E,F,G,H | 3.6295362 |
| *Prss56* | 12 | M | D,E,F,G,H,I | 3.6191376 |
| $C1qtnf5^{tm1.1(KOMP)Vlcg}$ | 4 | M | D,E,F,G,H,I,J | 3.6050767 |
| *Adipor1* | 8 | M | E,F,G,H,I,J,K | 3.5902411 |
| *Adipor1* | 8 | F | F,G,H,I,J,K | 3.5875206 |
| *Prss56* | 12 | F | G,H,I,J,K,L | 3.5798547 |
| *Mfrp* | 12 | M | G,H,I,J,K,L | 3.5793219 |
| *Mfrp* | 12 | F | I,J,K,L | 3.5634515 |
| *B6J* | 4 | M | J,K,L | 3.5624256 |
| $C1qtnf5^{tm1.1(KOMP)Vlcg}$ | 4 | F | H,I,J,K,L,M,N,O | 3.5489981 |
| *B6J* | 4 | F | J,K,L,M,N | 3.5434722 |
| *Mfrp* | 8 | F | L,M,N,O | 3.5297461 |
| *Prss56* | 8 | F | K,L,M,N,O,P | 3.5255357 |
| *Mfrp* | 8 | M | L,M,N,O,P | 3.5216087 |
| *Prss56* | 8 | M | M,N,O,P | 3.5200877 |
| *Adipor1* | 4 | M | N,O,P,Q | 3.4904908 |
| *Adipor1* | 4 | F | O,P,Q | 3.4869922 |
| *Prss56* | 4 | M | P,Q | 3.4635476 |
| *Prss56* | 4 | F | P,Q | 3.4624926 |
| *Mfrp* | 4 | F | P,Q | 3.4574057 |
| *Mfrp* | 4 | M | Q | 3.4519363 |
| *Adipor1* | 1 | M | R,S | 3.2750911 |
| *B6J* | 1 | M | R | 3.2669944 |
| *Adipor1* | 1 | F | R,S,T | 3.2481718 |
| *B6J* | 1 | F | R,S,T | 3.2372444 |
| *Prss56* | 1 | M | R,S,T | 3.2350815 |
| $C1qtnf5^{tm1.1(KOMP)Vlcg}$ | 1 | M | R,S,T | 3.2291575 |
| $C1qtnf5^{tm1.1(KOMP)Vlcg}$ | 1 | F | R,S,T | 3.2144605 |
| *Mfrp* | 1 | F | S,T | 3.2078336 |
| *Prss56* | 1 | F | T | 3.2039729 |
| *Mfrp* | 1 | M | T | 3.2008170 |

**Table ST3:** Least squares means (LSMeans) of strains used in this study. Strains not connected by the same letter are significantly different from Tukey's HSD post-hoc test.

| Tukey's multiple comparison | Adjusted p-value | Summary |
|---|---|---|
| **Age (1 month)** | | |
| C57BL/6J vs. *Mfrp*$^{rd6}$ | 0.9195 | ns |
| C57BL/6J vs. *Prss56*$^{glcr4}$ | 0.6371 | ns |
| C57BL/6J vs. *Adipor1*$^{tm1Dgen}$ | 0.6983 | ns |
| C57BL/6J vs. *C1qtnf5*$^{tm1.1(KOMP)Vlcg}$ | 0.0565 | ns |
| *Mfrp*$^{rd6}$ vs. *Prss56*$^{glcr4}$ | 0.9690 | ns |
| *Mfrp*$^{rd6}$ vs. *Adipor1*$^{tm1Dgen}$ | 0.9684 | ns |
| *Mfrp*$^{rd6}$ vs. *C1qtnf5*$^{tm1.1(KOMP)Vlcg}$ | 0.3000 | ns |
| *Prss56*$^{glcr4}$ vs. *Adipor1*$^{tm1Dgen}$ | >0.9999 | ns |
| *Prss56*$^{glcr4}$ vs. *C1qtnf5*$^{tm1.1(KOMP)Vlcg}$ | 0.7667 | ns |
| *Adipor1*$^{tm1Dgen}$ vs. *C1qtnf5*$^{tm1.1(KOMP)Vlcg}$ | 0.8587 | ns |
| **Age (4 month)** | | |
| C57BL/6J vs. *Mfrp*$^{rd6}$ | 0.1209 | ns |
| C57BL/6J vs. *Prss56*$^{glcr4}$ | 0.9075 | ns |
| C57BL/6J vs. *Adipor1*$^{tm1Dgen}$ | >0.9999 | ns |
| C57BL/6J vs. *C1qtnf5*$^{tm1.1(KOMP)Vlcg}$ | 0.9997 | ns |
| *Mfrp*$^{rd6}$ vs. *Prss56*$^{glcr4}$ | 0.0433 | * |
| *Mfrp*$^{rd6}$ vs. *Adipor1*$^{tm1Dgen}$ | 0.1448 | ns |
| *Mfrp*$^{rd6}$ vs. *C1qtnf5*$^{tm1.1(KOMP)Vlcg}$ | 0.4376 | ns |
| *Prss56*$^{glcr4}$ vs. *Adipor1*$^{tm1Dgen}$ | 0.8970 | ns |
| *Prss56*$^{glcr4}$ vs. *C1qtnf5*$^{tm1.1(KOMP)Vlcg}$ | 0.9385 | ns |
| *Adipor1*$^{tm1Dgen}$ vs. *C1qtnf5*$^{tm1.1(KOMP)Vlcg}$ | >0.9999 | ns |
| **Age (8 month)** | | |
| C57BL/6J vs. *Mfrp*$^{rd6}$ | 0.0326 | * |
| C57BL/6J vs. *Prss56*$^{glcr4}$ | 0.9991 | ns |
| C57BL/6J vs. *Adipor1*$^{tm1Dgen}$ | 0.9272 | ns |
| C57BL/6J vs. *C1qtnf5*$^{tm1.1(KOMP)Vlcg}$ | 0.8244 | ns |
| *Mfrp*$^{rd6}$ vs. *Prss56*$^{glcr4}$ | 0.3224 | ns |
| *Mfrp*$^{rd6}$ vs. *Adipor1*$^{tm1Dgen}$ | 0.0041 | ** |
| *Mfrp*$^{rd6}$ vs. *C1qtnf5*$^{tm1.1(KOMP)Vlcg}$ | 0.4563 | ns |
| *Prss56*$^{glcr4}$ vs. *Adipor1*$^{tm1Dgen}$ | 0.9205 | ns |
| *Prss56*$^{glcr4}$ vs. *C1qtnf5*$^{tm1.1(KOMP)Vlcg}$ | 0.9798 | ns |
| *Adipor1*$^{tm1Dgen}$ vs. *C1qtnf5*$^{tm1.1(KOMP)Vlcg}$ | 0.4105 | ns |
| **Age (12 month)** | | |
| C57BL/6J vs. *Mfrp*$^{rd6}$ | 0.9641 | ns |
| C57BL/6J vs. *Prss56*$^{glcr4}$ | 0.9218 | ns |
| C57BL/6J vs. *Adipor1*$^{tm1Dgen}$ | 0.9666 | ns |
| C57BL/6J vs. *C1qtnf5*$^{tm1.1(KOMP)Vlcg}$ | 0.6242 | ns |
| *Mfrp*$^{rd6}$ vs. *Prss56*$^{glcr4}$ | >0.9999 | ns |
| *Mfrp*$^{rd6}$ vs. *Adipor1*$^{tm1Dgen}$ | >0.9999 | ns |
| *Mfrp*$^{rd6}$ vs. *C1qtnf5*$^{tm1.1(KOMP)Vlcg}$ | 0.2936 | ns |
| *Prss56*$^{glcr4}$ vs. *Adipor1*$^{tm1Dgen}$ | 0.9999 | ns |
| *Prss56*$^{glcr4}$ vs. *C1qtnf5*$^{tm1.1(KOMP)Vlcg}$ | 0.2149 | ns |
| *Adipor1*$^{tm1Dgen}$ vs. *C1qtnf5*$^{tm1.1(KOMP)Vlcg}$ | 0.2915 | ns |

**Table ST4:** Adjusted *p*-values obtained from Tukey's multiple comparison test for CT comparison between different mouse strains, at four different ages.

| Tukey's multiple comparison | Adjusted p-value | Summary |
|---|---|---|
| **Age (1 month)** | | |
| C57BL/6J vs. *Mfrp*$^{rd6}$ | 0.0623 | ns |
| C57BL/6J vs. *Prss56*$^{glcr4}$ | <0.0001 | **** |
| C57BL/6J vs. *Adipor1*$^{tm1Dgen}$ | 0.0096 | ** |
| C57BL/6J vs. *C1qtnf5*$^{tm1.1(KOMP)Vlcg}$ | 0.0005 | *** |
| *Mfrp*$^{rd6}$ vs. *Prss56*$^{glcr4}$ | 0.0172 | * |
| *Mfrp*$^{rd6}$ vs. *Adipor1*$^{tm1Dgen}$ | 0.6576 | ns |
| *Mfrp*$^{rd6}$ vs. *C1qtnf5*$^{tm1.1(KOMP)Vlcg}$ | <0.0001 | **** |
| *Prss56*$^{glcr4}$ vs. *Adipor1*$^{tm1Dgen}$ | 0.6716 | ns |
| *Prss56*$^{glcr4}$ vs. *C1qtnf5*$^{tm1.1(KOMP)Vlcg}$ | <0.0001 | **** |
| *Adipor1*$^{tm1Dgen}$ vs. *C1qtnf5*$^{tm1.1(KOMP)Vlcg}$ | <0.0001 | **** |
| **Age (4 month)** | | |
| C57BL/6J vs. *Mfrp*$^{rd6}$ | <0.0001 | **** |
| C57BL/6J vs. *Prss56*$^{glcr4}$ | <0.0001 | **** |
| C57BL/6J vs. *Adipor1*$^{tm1Dgen}$ | 0.0062 | ** |
| C57BL/6J vs. *C1qtnf5*$^{tm1.1(KOMP)Vlcg}$ | 0.1713 | ns |
| *Mfrp*$^{rd6}$ vs. *Prss56*$^{glcr4}$ | 0.1009 | ns |
| *Mfrp*$^{rd6}$ vs. *Adipor1*$^{tm1Dgen}$ | <0.0001 | **** |
| *Mfrp*$^{rd6}$ vs. *C1qtnf5*$^{tm1.1(KOMP)Vlcg}$ | 0.0033 | ** |
| *Prss56*$^{glcr4}$ vs. *Adipor1*$^{tm1Dgen}$ | <0.0001 | **** |
| *Prss56*$^{glcr4}$ vs. *C1qtnf5*$^{tm1.1(KOMP)Vlcg}$ | 0.0001 | *** |
| *Adipor1*$^{tm1Dgen}$ vs. *C1qtnf5*$^{tm1.1(KOMP)Vlcg}$ | 0.9948 | ns |
| **Age (8 month)** | | |
| C57BL/6J vs. *Mfrp*$^{rd6}$ | <0.0001 | **** |
| C57BL/6J vs. *Prss56*$^{glcr4}$ | <0.0001 | **** |
| C57BL/6J vs. *Adipor1*$^{tm1Dgen}$ | 0.0230 | * |
| C57BL/6J vs. *C1qtnf5*$^{tm1.1(KOMP)Vlcg}$ | 0.9033 | ns |
| *Mfrp*$^{rd6}$ vs. *Prss56*$^{glcr4}$ | 0.8964 | ns |
| *Mfrp*$^{rd6}$ vs. *Adipor1*$^{tm1Dgen}$ | 0.0313 | * |
| *Mfrp*$^{rd6}$ vs. *C1qtnf5*$^{tm1.1(KOMP)Vlcg}$ | <0.0001 | **** |
| *Prss56*$^{glcr4}$ vs. *Adipor1*$^{tm1Dgen}$ | 0.0082 | ** |
| *Prss56*$^{glcr4}$ vs. *C1qtnf5*$^{tm1.1(KOMP)Vlcg}$ | <0.0001 | **** |
| *Adipor1*$^{tm1Dgen}$ vs. *C1qtnf5*$^{tm1.1(KOMP)Vlcg}$ | 0.0070 | ** |
| **Age (12 month)** | | |
| C57BL/6J vs. *Mfrp*$^{rd6}$ | <0.0001 | **** |
| C57BL/6J vs. *Prss56*$^{glcr4}$ | <0.0001 | **** |
| C57BL/6J vs. *Adipor1*$^{tm1Dgen}$ | <0.0001 | **** |
| C57BL/6J vs. *C1qtnf5*$^{tm1.1(KOMP)Vlcg}$ | 0.3770 | ns |
| *Mfrp*$^{rd6}$ vs. *Prss56*$^{glcr4}$ | 0.8008 | ns |
| *Mfrp*$^{rd6}$ vs. *Adipor1*$^{tm1Dgen}$ | 0.0198 | * |
| *Mfrp*$^{rd6}$ vs. *C1qtnf5*$^{tm1.1(KOMP)Vlcg}$ | <0.0001 | **** |
| *Prss56*$^{glcr4}$ vs. *Adipor1*$^{tm1Dgen}$ | 0.0003 | *** |
| *Prss56*$^{glcr4}$ vs. *C1qtnf5*$^{tm1.1(KOMP)Vlcg}$ | <0.0001 | **** |
| *Adipor1*$^{tm1Dgen}$ vs. *C1qtnf5*$^{tm1.1(KOMP)Vlcg}$ | <0.0001 | **** |

**Table ST5:** Adjusted *p*-values obtained from Tukey's multiple comparison test for ACD comparison between different mouse strains, at four different ages.

| Tukey's multiple comparison | Adjusted p-value | Summary |
|---|---|---|
| **Age (1 month)** | | |
| C57BL/6J vs. $Mfrp^{rd6}$ | 0.9508 | ns |
| C57BL/6J vs. $Prss56^{glcr4}$ | 0.3358 | ns |
| C57BL/6J vs. $Adipor1^{tm1Dgen}$ | 0.5437 | ns |
| C57BL/6J vs. $C1qtnf5^{tm1.1(KOMP)Vlcg}$ | 0.0104 | * |
| $Mfrp^{rd6}$ vs. $Prss56^{glcr4}$ | 0.9709 | ns |
| $Mfrp^{rd6}$ vs. $Adipor1^{tm1Dgen}$ | 0.9988 | ns |
| $Mfrp^{rd6}$ vs. $C1qtnf5^{tm1.1(KOMP)Vlcg}$ | 0.0217 | * |
| $Prss56^{glcr4}$ vs. $Adipor1^{tm1Dgen}$ | 0.9826 | ns |
| $Prss56^{glcr4}$ vs. $C1qtnf5^{tm1.1(KOMP)Vlcg}$ | 0.0003 | *** |
| $Adipor1^{tm1Dgen}$ vs. $C1qtnf5^{tm1.1(KOMP)Vlcg}$ | 0.0007 | *** |
| **Age (4 month)** | | |
| C57BL/6J vs. $Mfrp^{rd6}$ | 0.2787 | ns |
| C57BL/6J vs. $Prss56^{glcr4}$ | 0.7042 | ns |
| C57BL/6J vs. $Adipor1^{tm1Dgen}$ | 0.9999 | ns |
| C57BL/6J vs. $C1qtnf5^{tm1.1(KOMP)Vlcg}$ | 0.9966 | ns |
| $Mfrp^{rd6}$ vs. $Prss56^{glcr4}$ | 0.0321 | * |
| $Mfrp^{rd6}$ vs. $Adipor1^{tm1Dgen}$ | 0.2291 | ns |
| $Mfrp^{rd6}$ vs. $C1qtnf5^{tm1.1(KOMP)Vlcg}$ | 0.4103 | ns |
| $Prss56^{glcr4}$ vs. $Adipor1^{tm1Dgen}$ | 0.7901 | ns |
| $Prss56^{glcr4}$ vs. $C1qtnf5^{tm1.1(KOMP)Vlcg}$ | 0.9613 | ns |
| $Adipor1^{tm1Dgen}$ vs. $C1qtnf5^{tm1.1(KOMP)Vlcg}$ | 0.9993 | ns |
| **Age (8 month)** | | |
| C57BL/6J vs. $Mfrp^{rd6}$ | >0.9999 | ns |
| C57BL/6J vs. $Prss56^{glcr4}$ | 0.0086 | ** |
| C57BL/6J vs. $Adipor1^{tm1Dgen}$ | 0.0953 | ns |
| C57BL/6J vs. $C1qtnf5^{tm1.1(KOMP)Vlcg}$ | 0.3927 | ns |
| $Mfrp^{rd6}$ vs. $Prss56^{glcr4}$ | 0.0025 | ** |
| $Mfrp^{rd6}$ vs. $Adipor1^{tm1Dgen}$ | 0.0398 | * |
| $Mfrp^{rd6}$ vs. $C1qtnf5^{tm1.1(KOMP)Vlcg}$ | 0.3749 | ns |
| $Prss56^{glcr4}$ vs. $Adipor1^{tm1Dgen}$ | 0.4305 | ns |
| $Prss56^{glcr4}$ vs. $C1qtnf5^{tm1.1(KOMP)Vlcg}$ | <0.0001 | **** |
| $Adipor1^{tm1Dgen}$ vs. $C1qtnf5^{tm1.1(KOMP)Vlcg}$ | 0.0008 | *** |
| **Age (12 month)** | | |
| C57BL/6J vs. $Mfrp^{rd6}$ | 0.2317 | ns |
| C57BL/6J vs. $Prss56^{glcr4}$ | <0.0001 | **** |
| C57BL/6J vs. $Adipor1^{tm1Dgen}$ | 0.0017 | ** |
| C57BL/6J vs. $C1qtnf5^{tm1.1(KOMP)Vlcg}$ | 0.5268 | ns |
| $Mfrp^{rd6}$ vs. $Prss56^{glcr4}$ | 0.0798 | ns |
| $Mfrp^{rd6}$ vs. $Adipor1^{tm1Dgen}$ | 0.5069 | ns |
| $Mfrp^{rd6}$ vs. $C1qtnf5^{tm1.1(KOMP)Vlcg}$ | 0.9484 | ns |
| $Prss56^{glcr4}$ vs. $Adipor1^{tm1Dgen}$ | 0.7051 | ns |
| $Prss56^{glcr4}$ vs. $C1qtnf5^{tm1.1(KOMP)Vlcg}$ | 0.0060 | ** |
| $Adipor1^{tm1Dgen}$ vs. $C1qtnf5^{tm1.1(KOMP)Vlcg}$ | 0.0905 | **** |

**Table ST6:** Adjusted *p*-values obtained from Tukey's multiple comparison test for LT comparison between different mouse strains, at four different ages.

| Tukey's multiple comparison | Adjusted p-value | Summary |
|---|---|---|
| **Age (1 month)** | | |
| C57BL/6J vs. *Mfrp*$^{rd6}$ | <0.0001 | **** |
| C57BL/6J vs. *Prss56*$^{glcr4}$ | 0.0701 | ns |
| C57BL/6J vs. *Adipor1*$^{tm1Dgen}$ | <0.0001 | **** |
| C57BL/6J vs. *C1qtnf5*$^{tm1.1(KOMP)Vlcg}$ | 0.0277 | * |
| *Mfrp*$^{rd6}$ vs. *Prss56*$^{glcr4}$ | <0.0001 | **** |
| *Mfrp*$^{rd6}$ vs. *Adipor1*$^{tm1Dgen}$ | 0.6086 | ns |
| *Mfrp*$^{rd6}$ vs. *C1qtnf5*$^{tm1.1(KOMP)Vlcg}$ | <0.0001 | **** |
| *Prss56*$^{glcr4}$ vs. *Adipor1*$^{tm1Dgen}$ | <0.0001 | **** |
| *Prss56*$^{glcr4}$ vs. *C1qtnf5*$^{tm1.1(KOMP)Vlcg}$ | <0.0001 | **** |
| *Adipor1*$^{tm1Dgen}$ vs. *C1qtnf5*$^{tm1.1(KOMP)Vlcg}$ | 0.0025 | ** |
| **Age (4 month)** | | |
| C57BL/6J vs. *Mfrp*$^{rd6}$ | <0.0001 | **** |
| C57BL/6J vs. *Prss56*$^{glcr4}$ | <0.0001 | **** |
| C57BL/6J vs. *Adipor1*$^{tm1Dgen}$ | <0.0001 | **** |
| C57BL/6J vs. *C1qtnf5*$^{tm1.1(KOMP)Vlcg}$ | <0.0001 | **** |
| *Mfrp*$^{rd6}$ vs. *Prss56*$^{glcr4}$ | <0.0001 | **** |
| *Mfrp*$^{rd6}$ vs. *Adipor1*$^{tm1Dgen}$ | 0.0148 | * |
| *Mfrp*$^{rd6}$ vs. *C1qtnf5*$^{tm1.1(KOMP)Vlcg}$ | <0.0001 | **** |
| *Prss56*$^{glcr4}$ vs. *Adipor1*$^{tm1Dgen}$ | <0.0001 | **** |
| *Prss56*$^{glcr4}$ vs. *C1qtnf5*$^{tm1.1(KOMP)Vlcg}$ | <0.0001 | **** |
| *Adipor1*$^{tm1Dgen}$ vs. *C1qtnf5*$^{tm1.1(KOMP)Vlcg}$ | <0.0001 | **** |
| **Age (8 month)** | | |
| C57BL/6J vs. *Mfrp*$^{rd6}$ | <0.0001 | **** |
| C57BL/6J vs. *Prss56*$^{glcr4}$ | <0.0001 | **** |
| C57BL/6J vs. *Adipor1*$^{tm1Dgen}$ | <0.0001 | **** |
| C57BL/6J vs. *C1qtnf5*$^{tm1.1(KOMP)Vlcg}$ | 0.1142 | ns |
| *Mfrp*$^{rd6}$ vs. *Prss56*$^{glcr4}$ | <0.0001 | **** |
| *Mfrp*$^{rd6}$ vs. *Adipor1*$^{tm1Dgen}$ | <0.0001 | **** |
| *Mfrp*$^{rd6}$ vs. *C1qtnf5*$^{tm1.1(KOMP)Vlcg}$ | <0.0001 | **** |
| *Prss56*$^{glcr4}$ vs. *Adipor1*$^{tm1Dgen}$ | <0.0001 | **** |
| *Prss56*$^{glcr4}$ vs. *C1qtnf5*$^{tm1.1(KOMP)Vlcg}$ | <0.0001 | **** |
| *Adipor1*$^{tm1Dgen}$ vs. *C1qtnf5*$^{tm1.1(KOMP)Vlcg}$ | <0.0001 | **** |
| **Age (12 month)** | | |
| C57BL/6J vs. *Mfrp*$^{rd6}$ | <0.0001 | **** |
| C57BL/6J vs. *Prss56*$^{glcr4}$ | 0.2878 | ns |
| C57BL/6J vs. *Adipor1*$^{tm1Dgen}$ | <0.0001 | **** |
| C57BL/6J vs. *C1qtnf5*$^{tm1.1(KOMP)Vlcg}$ | <0.0001 | **** |
| *Mfrp*$^{rd6}$ vs. *Prss56*$^{glcr4}$ | <0.0001 | **** |
| *Mfrp*$^{rd6}$ vs. *Adipor1*$^{tm1Dgen}$ | <0.0001 | **** |
| *Mfrp*$^{rd6}$ vs. *C1qtnf5*$^{tm1.1(KOMP)Vlcg}$ | <0.0001 | **** |
| *Prss56*$^{glcr4}$ vs. *Adipor1*$^{tm1Dgen}$ | <0.0001 | **** |
| *Prss56*$^{glcr4}$ vs. *C1qtnf5*$^{tm1.1(KOMP)Vlcg}$ | <0.0001 | **** |
| *Adipor1*$^{tm1Dgen}$ vs. *C1qtnf5*$^{tm1.1(KOMP)Vlcg}$ | <0.0001 | **** |

**Table ST7:** Adjusted *p*-values obtained from Tukey's multiple comparison test for ONL T comparison between different mouse strains, at four different ages.

| Tukey's multiple comparison | Adjusted p-value | Summary |
|---|---|---|
| **Age (1 month)** | | |
| C57BL/6J vs. *Mfrp*$^{rd6}$ | <0.0001 | **** |
| C57BL/6J vs. *Prss56*$^{glcr4}$ | <0.0001 | **** |
| C57BL/6J vs. *Adipor1*$^{tm1Dgen}$ | <0.0001 | **** |
| C57BL/6J vs. *C1qtnf5*$^{tm1.1(KOMP)Vlcg}$ | 0.0078 | ** |
| *Mfrp*$^{rd6}$ vs. *Prss56*$^{glcr4}$ | <0.0001 | **** |
| *Mfrp*$^{rd6}$ vs. *Adipor1*$^{tm1Dgen}$ | <0.0001 | **** |
| *Mfrp*$^{rd6}$ vs. *C1qtnf5*$^{tm1.1(KOMP)Vlcg}$ | <0.0001 | **** |
| *Prss56*$^{glcr4}$ vs. *Adipor1*$^{tm1Dgen}$ | <0.0001 | **** |
| *Prss56*$^{glcr4}$ vs. *C1qtnf5*$^{tm1.1(KOMP)Vlcg}$ | <0.0001 | **** |
| *Adipor1*$^{tm1Dgen}$ vs. *C1qtnf5*$^{tm1.1(KOMP)Vlcg}$ | <0.0001 | **** |
| **Age (4 month)** | | |
| C57BL/6J vs. *Mfrp*$^{rd6}$ | <0.0001 | **** |
| C57BL/6J vs. *Prss56*$^{glcr4}$ | <0.0001 | **** |
| C57BL/6J vs. *Adipor1*$^{tm1Dgen}$ | <0.0001 | **** |
| C57BL/6J vs. *C1qtnf5*$^{tm1.1(KOMP)Vlcg}$ | 0.4116 | ns |
| *Mfrp*$^{rd6}$ vs. *Prss56*$^{glcr4}$ | <0.0001 | **** |
| *Mfrp*$^{rd6}$ vs. *Adipor1*$^{tm1Dgen}$ | <0.0001 | **** |
| *Mfrp*$^{rd6}$ vs. *C1qtnf5*$^{tm1.1(KOMP)Vlcg}$ | <0.0001 | **** |
| *Prss56*$^{glcr4}$ vs. *Adipor1*$^{tm1Dgen}$ | <0.0001 | **** |
| *Prss56*$^{glcr4}$ vs. *C1qtnf5*$^{tm1.1(KOMP)Vlcg}$ | <0.0001 | **** |
| *Adipor1*$^{tm1Dgen}$ vs. *C1qtnf5*$^{tm1.1(KOMP)Vlcg}$ | <0.0001 | **** |
| **Age (8 month)** | | |
| C57BL/6J vs. *Mfrp*$^{rd6}$ | <0.0001 | **** |
| C57BL/6J vs. *Prss56*$^{glcr4}$ | <0.0001 | **** |
| C57BL/6J vs. *Adipor1*$^{tm1Dgen}$ | <0.0001 | **** |
| C57BL/6J vs. *C1qtnf5*$^{tm1.1(KOMP)Vlcg}$ | 0.0424 | * |
| *Mfrp*$^{rd6}$ vs. *Prss56*$^{glcr4}$ | <0.0001 | **** |
| *Mfrp*$^{rd6}$ vs. *Adipor1*$^{tm1Dgen}$ | <0.0001 | **** |
| *Mfrp*$^{rd6}$ vs. *C1qtnf5*$^{tm1.1(KOMP)Vlcg}$ | <0.0001 | **** |
| *Prss56*$^{glcr4}$ vs. *Adipor1*$^{tm1Dgen}$ | <0.0001 | **** |
| *Prss56*$^{glcr4}$ vs. *C1qtnf5*$^{tm1.1(KOMP)Vlcg}$ | <0.0001 | **** |
| *Adipor1*$^{tm1Dgen}$ vs. *C1qtnf5*$^{tm1.1(KOMP)Vlcg}$ | <0.0001 | **** |
| **Age (12 month)** | | |
| C57BL/6J vs. *Mfrp*$^{rd6}$ | <0.0001 | **** |
| C57BL/6J vs. *Prss56*$^{glcr4}$ | <0.0001 | **** |
| C57BL/6J vs. *Adipor1*$^{tm1Dgen}$ | <0.0001 | **** |
| C57BL/6J vs. *C1qtnf5*$^{tm1.1(KOMP)Vlcg}$ | 0.0911 | ns |
| *Mfrp*$^{rd6}$ vs. *Prss56*$^{glcr4}$ | <0.0001 | **** |
| *Mfrp*$^{rd6}$ vs. *Adipor1*$^{tm1Dgen}$ | <0.0001 | **** |
| *Mfrp*$^{rd6}$ vs. *C1qtnf5*$^{tm1.1(KOMP)Vlcg}$ | <0.0001 | **** |
| *Prss56*$^{glcr4}$ vs. *Adipor1*$^{tm1Dgen}$ | <0.0001 | **** |
| *Prss56*$^{glcr4}$ vs. *C1qtnf5*$^{tm1.1(KOMP)Vlcg}$ | <0.0001 | **** |
| *Adipor1*$^{tm1Dgen}$ vs. *C1qtnf5*$^{tm1.1(KOMP)Vlcg}$ | <0.0001 | **** |

**Table ST8:** Adjusted *p*-values obtained from Tukey's multiple comparison test for RT comparison between different mouse strains, at four different ages.

| Tukey's multiple comparison | Adjusted p-value | Summary |
|---|---|---|
| **Age (1 month)** | | |
| C57BL/6J vs. *Mfrp*$^{rd6}$ | <0.0001 | **** |
| C57BL/6J vs. *Prss56*$^{glcr4}$ | <0.0001 | **** |
| C57BL/6J vs. *Adipor1*$^{tm1Dgen}$ | 0.3607 | ns |
| C57BL/6J vs. *C1qtnf5*$^{tm1.1(KOMP)Vlcg}$ | 0.2284 | ns |
| *Mfrp*$^{rd6}$ vs. *Prss56*$^{glcr4}$ | <0.0001 | **** |
| *Mfrp*$^{rd6}$ vs. *Adipor1*$^{tm1Dgen}$ | <0.0001 | **** |
| *Mfrp*$^{rd6}$ vs. *C1qtnf5*$^{tm1.1(KOMP)Vlcg}$ | <0.0001 | **** |
| *Prss56*$^{glcr4}$ vs. *Adipor1*$^{tm1Dgen}$ | <0.0001 | **** |
| *Prss56*$^{glcr4}$ vs. *C1qtnf5*$^{tm1.1(KOMP)Vlcg}$ | <0.0001 | **** |
| *Adipor1*$^{tm1Dgen}$ vs. *C1qtnf5*$^{tm1.1(KOMP)Vlcg}$ | >0.9999 | ns |
| **Age (4 month)** | | |
| C57BL/6J vs. *Mfrp*$^{rd6}$ | <0.0001 | **** |
| C57BL/6J vs. *Prss56*$^{glcr4}$ | <0.0001 | **** |
| C57BL/6J vs. *Adipor1*$^{tm1Dgen}$ | 0.0005 | *** |
| C57BL/6J vs. *C1qtnf5*$^{tm1.1(KOMP)Vlcg}$ | 0.9986 | ns |
| *Mfrp*$^{rd6}$ vs. *Prss56*$^{glcr4}$ | <0.0001 | **** |
| *Mfrp*$^{rd6}$ vs. *Adipor1*$^{tm1Dgen}$ | <0.0001 | **** |
| *Mfrp*$^{rd6}$ vs. *C1qtnf5*$^{tm1.1(KOMP)Vlcg}$ | <0.0001 | **** |
| *Prss56*$^{glcr4}$ vs. *Adipor1*$^{tm1Dgen}$ | <0.0001 | **** |
| *Prss56*$^{glcr4}$ vs. *C1qtnf5*$^{tm1.1(KOMP)Vlcg}$ | <0.0001 | **** |
| *Adipor1*$^{tm1Dgen}$ vs. *C1qtnf5*$^{tm1.1(KOMP)Vlcg}$ | 0.0005 | *** |
| **Age (8 month)** | | |
| C57BL/6J vs. *Mfrp*$^{rd6}$ | <0.0001 | **** |
| C57BL/6J vs. *Prss56*$^{glcr4}$ | <0.0001 | **** |
| C57BL/6J vs. *Adipor1*$^{tm1Dgen}$ | <0.0001 | **** |
| C57BL/6J vs. *C1qtnf5*$^{tm1.1(KOMP)Vlcg}$ | 0.0022 | ** |
| *Mfrp*$^{rd6}$ vs. *Prss56*$^{glcr4}$ | <0.0001 | **** |
| *Mfrp*$^{rd6}$ vs. *Adipor1*$^{tm1Dgen}$ | <0.0001 | **** |
| *Mfrp*$^{rd6}$ vs. *C1qtnf5*$^{tm1.1(KOMP)Vlcg}$ | <0.0001 | **** |
| *Prss56*$^{glcr4}$ vs. *Adipor1*$^{tm1Dgen}$ | <0.0001 | **** |
| *Prss56*$^{glcr4}$ vs. *C1qtnf5*$^{tm1.1(KOMP)Vlcg}$ | <0.0001 | **** |
| *Adipor1*$^{tm1Dgen}$ vs. *C1qtnf5*$^{tm1.1(KOMP)Vlcg}$ | <0.0001 | **** |
| **Age (12 month)** | | |
| C57BL/6J vs. *Mfrp*$^{rd6}$ | <0.0001 | **** |
| C57BL/6J vs. *Prss56*$^{glcr4}$ | <0.0001 | **** |
| C57BL/6J vs. *Adipor1*$^{tm1Dgen}$ | 0.3209 | ns |
| C57BL/6J vs. *C1qtnf5*$^{tm1.1(KOMP)Vlcg}$ | <0.0001 | **** |
| *Mfrp*$^{rd6}$ vs. *Prss56*$^{glcr4}$ | <0.0001 | **** |
| *Mfrp*$^{rd6}$ vs. *Adipor1*$^{tm1Dgen}$ | <0.0001 | **** |
| *Mfrp*$^{rd6}$ vs. *C1qtnf5*$^{tm1.1(KOMP)Vlcg}$ | <0.0001 | **** |
| *Prss56*$^{glcr4}$ vs. *Adipor1*$^{tm1Dgen}$ | <0.0001 | **** |
| *Prss56*$^{glcr4}$ vs. *C1qtnf5*$^{tm1.1(KOMP)Vlcg}$ | <0.0001 | **** |
| *Adipor1*$^{tm1Dgen}$ vs. *C1qtnf5*$^{tm1.1(KOMP)Vlcg}$ | <0.0001 | **** |

**Table ST9:** Adjusted *p*-values obtained from Tukey's multiple comparison test for VCD comparison between different mouse strains, at four different ages.

| Tukey's multiple comparison | Adjusted p-value | Summary |
|---|---|---|
| **Age (1 month)** | | |
| C57BL/6J vs. *Mfrp*$^{rd6}$ | <0.0001 | **** |
| C57BL/6J vs. *Prss56*$^{glcr4}$ | <0.0001 | **** |
| C57BL/6J vs. *Adipor1*$^{tm1Dgen}$ | 0.0162 | * |
| C57BL/6J vs. *C1qtnf5*$^{tm1.1(KOMP)Vlcg}$ | 0.0232 | * |
| *Mfrp*$^{rd6}$ vs. *Prss56*$^{glcr4}$ | 0.7749 | ns |
| *Mfrp*$^{rd6}$ vs. *Adipor1*$^{tm1Dgen}$ | 0.0004 | *** |
| *Mfrp*$^{rd6}$ vs. *C1qtnf5*$^{tm1.1(KOMP)Vlcg}$ | <0.0001 | **** |
| *Prss56*$^{glcr4}$ vs. *Adipor1*$^{tm1Dgen}$ | <0.0001 | **** |
| *Prss56*$^{glcr4}$ vs. *C1qtnf5*$^{tm1.1(KOMP)Vlcg}$ | <0.0001 | **** |
| *Adipor1*$^{tm1Dgen}$ vs. *C1qtnf5*$^{tm1.1(KOMP)Vlcg}$ | <0.0001 | **** |
| **Age (4 month)** | | |
| C57BL/6J vs. *Mfrp*$^{rd6}$ | <0.0001 | **** |
| C57BL/6J vs. *Prss56*$^{glcr4}$ | <0.0001 | **** |
| C57BL/6J vs. *Adipor1*$^{tm1Dgen}$ | <0.0001 | **** |
| C57BL/6J vs. *C1qtnf5*$^{tm1.1(KOMP)Vlcg}$ | 0.7983 | ns |
| *Mfrp*$^{rd6}$ vs. *Prss56*$^{glcr4}$ | 0.2517 | ns |
| *Mfrp*$^{rd6}$ vs. *Adipor1*$^{tm1Dgen}$ | <0.0001 | **** |
| *Mfrp*$^{rd6}$ vs. *C1qtnf5*$^{tm1.1(KOMP)Vlcg}$ | <0.0001 | **** |
| *Prss56*$^{glcr4}$ vs. *Adipor1*$^{tm1Dgen}$ | <0.0001 | **** |
| *Prss56*$^{glcr4}$ vs. *C1qtnf5*$^{tm1.1(KOMP)Vlcg}$ | <0.0001 | **** |
| *Adipor1*$^{tm1Dgen}$ vs. *C1qtnf5*$^{tm1.1(KOMP)Vlcg}$ | <0.0001 | **** |
| **Age (8 month)** | | |
| C57BL/6J vs. *Mfrp*$^{rd6}$ | <0.0001 | **** |
| C57BL/6J vs. *Prss56*$^{glcr4}$ | <0.0001 | **** |
| C57BL/6J vs. *Adipor1*$^{tm1Dgen}$ | <0.0001 | **** |
| C57BL/6J vs. *C1qtnf5*$^{tm1.1(KOMP)Vlcg}$ | 0.0005 | *** |
| *Mfrp*$^{rd6}$ vs. *Prss56*$^{glcr4}$ | <0.0001 | **** |
| *Mfrp*$^{rd6}$ vs. *Adipor1*$^{tm1Dgen}$ | <0.0001 | **** |
| *Mfrp*$^{rd6}$ vs. *C1qtnf5*$^{tm1.1(KOMP)Vlcg}$ | <0.0001 | **** |
| *Prss56*$^{glcr4}$ vs. *Adipor1*$^{tm1Dgen}$ | <0.0001 | **** |
| *Prss56*$^{glcr4}$ vs. *C1qtnf5*$^{tm1.1(KOMP)Vlcg}$ | <0.0001 | **** |
| *Adipor1*$^{tm1Dgen}$ vs. *C1qtnf5*$^{tm1.1(KOMP)Vlcg}$ | <0.0001 | **** |
| **Age (12 month)** | | |
| C57BL/6J vs. *Mfrp*$^{rd6}$ | <0.0001 | **** |
| C57BL/6J vs. *Prss56*$^{glcr4}$ | <0.0001 | **** |
| C57BL/6J vs. *Adipor1*$^{tm1Dgen}$ | <0.0001 | **** |
| C57BL/6J vs. *C1qtnf5*$^{tm1.1(KOMP)Vlcg}$ | <0.0001 | **** |
| *Mfrp*$^{rd6}$ vs. *Prss56*$^{glcr4}$ | 0.4611 | ns |
| *Mfrp*$^{rd6}$ vs. *Adipor1*$^{tm1Dgen}$ | <0.0001 | **** |
| *Mfrp*$^{rd6}$ vs. *C1qtnf5*$^{tm1.1(KOMP)Vlcg}$ | <0.0001 | **** |
| *Prss56*$^{glcr4}$ vs. *Adipor1*$^{tm1Dgen}$ | <0.0001 | **** |
| *Prss56*$^{glcr4}$ vs. *C1qtnf5*$^{tm1.1(KOMP)Vlcg}$ | <0.0001 | **** |
| *Adipor1*$^{tm1Dgen}$ vs. *C1qtnf5*$^{tm1.1(KOMP)Vlcg}$ | <0.0001 | **** |

**Table ST10:** Adjusted *p*-values obtained from Tukey's multiple comparison test for PL comparison between different mouse strains, at four different ages.
